# Novel miRNA-inducing drugs enable differentiation of retinoic acid-resistant neuroblastoma cells

**DOI:** 10.1101/2024.06.05.597584

**Authors:** Lien D. Nguyen, Satyaki Sengupta, Kevin Cho, Alexander Floru, Rani E. George, Anna M. Krichevsky

**Affiliations:** Department of Neurology, Brigham and Women’s Hospital and Harvard Medical School, Boston, MA 02115, USA; Department of Pediatric Oncology, Dana-Farber Cancer Institute and Harvard Medical School, Boston, MA, 02115, USA

**Keywords:** Neuroblastoma, differentiation therapy, miR-124, MES, ADRN

## Abstract

Tumor cell heterogeneity in neuroblastoma, a pediatric cancer arising from neural crest-derived progenitor cells, poses a significant clinical challenge. In particular, unlike adrenergic (ADRN) neuroblastoma cells, mesenchymal (MES) cells are resistant to chemotherapy and retinoid therapy and thereby significantly contribute to relapses and treatment failures. Previous research suggested that overexpression or activation of miR-124, a neurogenic microRNA with tumor suppressor activity, can induce the differentiation of retinoic acid-resistant neuroblastoma cells. Leveraging our established screen for miRNA-modulatory small molecules, we validated PP121, a dual inhibitor of tyrosine and phosphoinositide kinases, as a robust inducer of miR-124. A combination of PP121 and BDNF-activating bufalin synergistically arrests proliferation, induces differentiation, and maintains the differentiated state of MES SK-N-AS cells for 8 weeks. RNA-seq and deconvolution analyses revealed a collapse of the ADRN core regulatory circuitry (CRC) and the emergence of novel CRCs associated with chromaffin cells and Schwann cell precursors. Using a similar protocol, we differentiated and maintained MES neuroblastoma GI-ME-N and SH-EP cell lines, as well as glioblastoma LN-229 and U-251 cell lines, for over 16 weeks. In conclusion, our novel protocol suggests a promising treatment for therapy-resistant cancers of the nervous system. Moreover, these long-lived, differentiated cells provide valuable models for studying mechanisms underlying differentiation, maturation, and senescence.

## Introduction

Neuroblastoma, a cancer originating from neural crest cells (NCCs) and the most common extracranial solid tumor of childhood, exhibits remarkable diversities in clinical presentations, treatment responses, prognosis, and tumor characteristics. Patients classified as low/immediate risk have an excellent 5-year survival rate of above 95%, whereas the survival rate for patients classified as high risk remains below 50% (*1, 2*). Cell lines derived from neuroblastoma tumors also display diversity in morphology, with earlier studies characterizing them as “N” cells, which have small, rounded cell bodies and numerous neurite-like processes, and “S” cells, which are larger, flattened, and highly substrate-adherent (*3–5*). Next-generation sequencing techniques have enabled the classification of neuroblastoma cells into two genetically distinct groups: the adrenergic (ADRN) cells, which are lineage-committed, and the more immature mesenchymal (MES) cells, which more closely resemble NCCs (*6, 7*). ADRN and MES cells can interconvert *in vitro* based on the switching of core regulatory circuitries (CRCs) (*7–9*). Notably, MES cells are generally resistant to standard treatments, including chemotherapy (*6, 7*), retinoic acid (RA) (*6*), inhibitors targeting anaplastic lymphoma kinase (ALK) (*10*), and therapeutic antibodies targeting disialoganglioside (GD2) (*11*). Furthermore, MES gene signature is enriched in relapsed tumors, suggesting that either ADRN cells convert to MES cells to escape treatments or that surviving MES cells repopulate new tumors (*12*). Overall, multiple evidence suggest an urgent need for treatments that effectively target MES neuroblastoma cells.

MicroRNAs (miRNAs) are small, noncoding RNAs that regulate gene expression by silencing mRNA targets. They play crucial roles in various cellular processes, including determining cell fate and identity (*13*). Particularly, miR-124 is one of the most enriched and abundant miRNAs in the nervous system and drives neuronal differentiation and neurite outgrowth (*14–18*). miR-124 is frequently downregulated in various types of cancers and is proposed as a key tumor suppressor targeting oncogenes and signaling pathways implicated in cancer progression (*19*). Multiple studies suggest that endogenous miR-124 is upregulated during RA-induced differentiation of neuroblastoma cells (*20–23*) and transfection of miR-124 mimics drives neuronal differentiation in neuroblastoma cell lines (*14, 24*). Therefore, we hypothesized that molecular pathways that upregulate miR-124 would promote the differentiation of neuroblastoma cells.

We previously developed a high-throughput screen for small molecule compounds that modulate neuronal miRNomes and established the first resource for miRNA-modulating drugs (*25, 26*). In this study, we leveraged this resource to test whether miR-124-inducing compounds inhibit proliferation and drive differentiation, particularly in RA-resistant MES cell lines. We validated PP121, a dual inhibitor of tyrosine and phosphoinositide kinases (*27*), as a robust activator of miR-124. In combination with bufalin, a drug that upregulates miR-132 and activates the brain-derived neurotrophic factor (BDNF) pathway (*25*), PP121 completely arrested cell proliferation, promoted cell elongation and sprouting of neurite-like processes, and sustained this differentiated state for 8 weeks. Transcriptomic analyses revealed a permanent downregulation of cell cycle pathways, an upregulation of immune-related and senescence pathways, and, unexpectedly, an enhanced MES signature and decreased ADRN signature. Similar to observations made in RA-induced differentiation models (*28*), PP121/bufalin treatment led to a collapse of the ADRN CRC. In contrast to RA models, the retino-sympathetic CRC that supports the RA-differentiated state was also downregulated following treatment. Instead, SK-N-AS cells differentiated towards a state that transcriptionally resembled chromaffin cells and Schwann cell precursors (SCPs) (*29*). We further showed that PP121 and bufalin similarly differentiate additional MES neuroblastoma cell lines including GI-ME-N and SH-EP, and the glioblastoma LN-229 and U-251 cell lines. In summary, our study presents a novel method for differentiating neuroblastoma cells and supports a potential new avenue for the treatment of MES neuroblastomas resistant to standard therapies.

## RESULTS

### PP121 upregulates miR-124, arrests proliferation, and promotes neurite-like processes in SK-N-AS cell

Although previous studies already implicated miR-124 in neuroblastoma differentiation (*14, 24*), we first aimed to confirm a causal relationship between miR-124 and differentiation in our cell line models. In humans, there are three miR-124 loci on chromosomes 2, 3, and 15 (*30*) that are processed into identical mature transcripts, with miR-124-3p being the major product. To determine whether exogenous miR-124 induces differentiation of MES neuroblastoma cells (*14, 24*), we transfected SK-N-AS cells with three miR-124-3p mimics corresponding to the three loci. We selected the SK-N-AS cell line because of its MES (*7*) or mixed identity (*6*) and its resistance to RA-induced neuronal differentiation (*31, 32*). 4 days after transfection, live cell imaging revealed the emergence of rare neuron-like cells characterized by narrow, rounded cell bodies and neurite-like processes in all miR-124 transfected conditions, which were absent in cells transfected with a non-targeted sequence (CTRL mimics) (Fig. 1A). miR-124-transfected cells also showed a ∼25% reduction in cell viability (Fig. 1B) and a ∼40% reduction in confluence (Fig. 1C). Transfection of MES GI-ME-N and SH-EP cells with miR-124 mimics similarly reduced cell viability and induced morphological changes (Sup. Fig.1A-D). These results suggested that exogenous miR-124 reduced MES cell proliferation and drove some cells toward acquiring a differentiated morphology.

**Figure 1:**
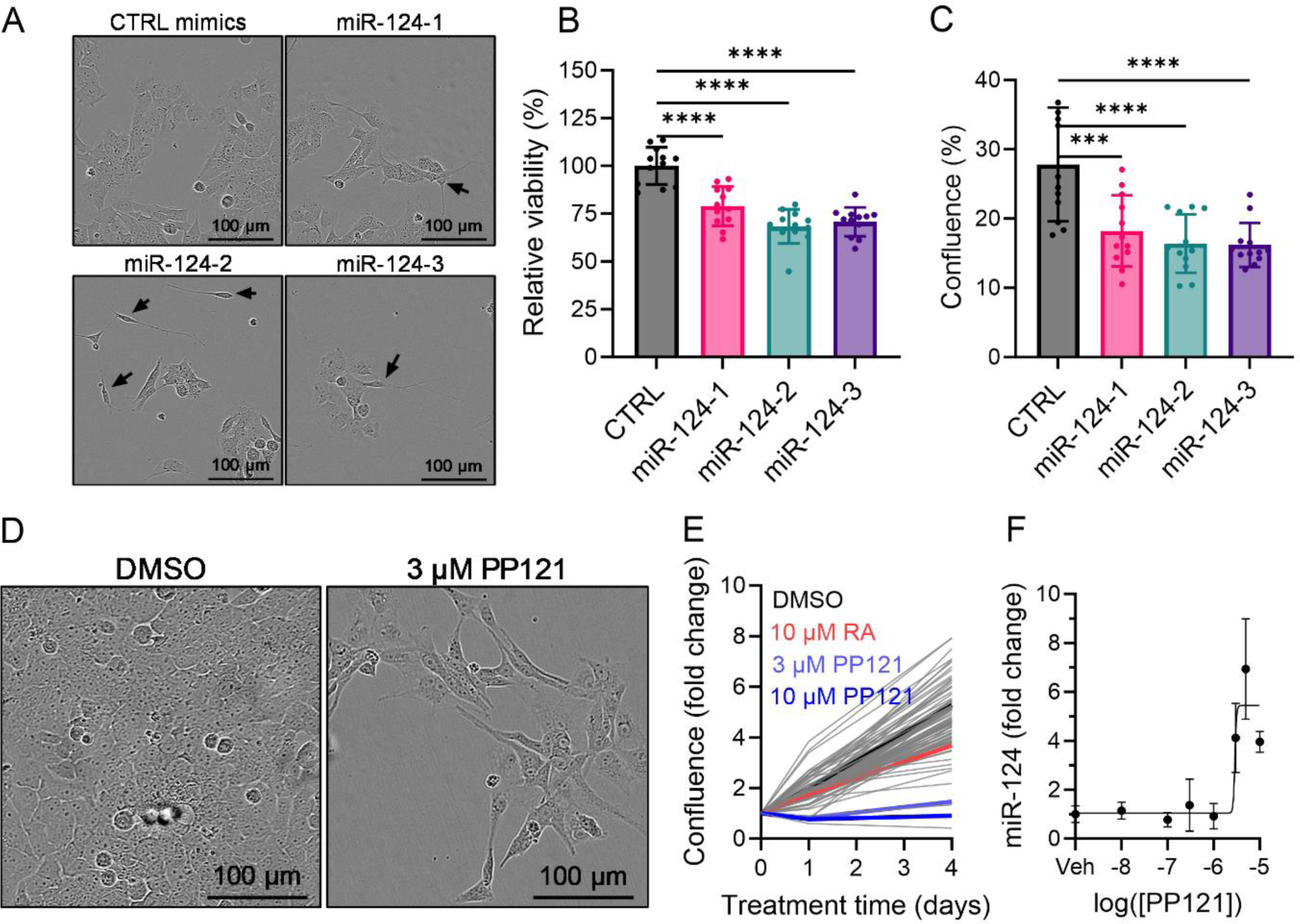
PP121 upregulates miR-124 and drives differentiation of SK-N-AS cells. **(A)** Transfection with miR-124 mimics for 4 days led some cells (black arrows) to acquire neuronal morphology and decreased **(B)** cell viability and **(C)** confluence growth (N=12, 1-way ANOVA, Dunnett’s multiple comparisons). **(D)** Representative images of SK-N-AS 4 days after treatment with DMSO and 3 µM PP121. **(E)** Normalized confluence growth over 4 days with treatments (N=1 per dose per drug, N=5 for DMSO). **(F)** 5 µM PP121 maximally upregulated miR-124 in SK-N-AS cells 7 days after treatment (N=4-6 per concentration). Error bars represent S.D.. Source data are provided in the source data file.

Based on these observations, we hypothesized that the activation of endogenous miR-124 in SK-N-AS cells with small molecule compounds would reduce proliferation and push cells toward a differentiated state. We proceeded to select 22 candidates with high miR-124-3p and −5p expression scores from our previous dataset (Supp. Fig. 2A, Supp. Table 1) (*25*). We treated SK-N-AS cells with these candidate compounds and RA at 4 doses (0.3, 1, 3, and 10 µM; N=1 well per dose) and determined morphological changes and cell growth rate using the IncuCyte® imaging system (Fig. 1D, Supp. Table 2). Notably, cells treated with 3 or 10 µM PP121, a multikinase inhibitor (*27*), stopped proliferation, elongated, and sprouted neurite-like processes (Fig. 1D, Supp. Fig. 2B). 10 µM RA, which was sufficient to induce differentiation in various ADRN cell lines (*28, 33*), slightly reduced growth (Fig. 1E), but did not differentiate SK-N-AS cells (Supp. Fig. 2B). Next, to determine whether PP121-mediated regulation of cell growth and morphology was associated with concomitant changes in miR-124 levels, we measured miR-124 abundance following treatment with increasing concentrations of the compound for 7 days. Indeed, PP121 upregulated miR-124 levels in a dose-dependent manner over 7 days, with maximal upregulation observed at 5 uM (Fig. 1F). In line with this observation, 5 uM PP121 differentiated SK-N-AS cells (Supp. Fig. 2C). However, the same concentration of PP121 completely killed ADRN-type Kelly and SK-N-SH cells (Supp. Fig. 2C). 5 uM PP121 did not appear to kill primary rat neurons, suggesting that PP121 primarily affected proliferating cells (Supp. Fig. 2C). Furthermore, the growth and morphological changes induced by PP121 was reversible, as the removal of PP121 by fresh medium exchange allowed SK-N-AS cells to resume proliferation (Supp. Fig. 2D).

### A novel protocol of PP121/bufalin induces long-term SK-N-AS differentiation

We observed that the morphology of SK-N-AS cells treated with PP121 resembled that of RA-differentiated neuroblastoma SH-SY-5Y cells. Therefore, we modified an established method for RA-induced differentiation of SH-SY-5Y cells (*33*) to see if we could further differentiate SK-N-AS cells. Briefly, in this protocol, SH-SY-5Y cells were differentiated through sequential culturing in growth medium (EMEM, 10% FBS), then differentiation medium #1 (EMEM, 2.5% FBS, 10 µM RA), differentiation medium #2 (EMEM, 1% FBS, 10 µM RA), and differentiation medium #3 (neurobasal, 1X B27, 20 mM KCl, 50 ng/mL BDNF, 2 mM db-cAMP, 10 µM RA) (*33*). In our initial modified protocol (Supp. Fig. 3A), 10 µM RA was replaced with 5 µM PP121. However, cells switched to neurobasal + 5 µM PP121 eventually clumped after several days, which was not ameliorated by adding BDNF, KCl, and db-cAMP, (Supp. Fig. 3B). We hypothesized that a broader activation of the BDN pathway might prevent cell clumping. As we and others previously found that cardiac glycosides activated the BDNF pathway in neurons (*25, 34*), we tested bufalin, a cardiac glycoside, to see if it could ameliorate cell clumping. Bufalin at a concentration of ≥100 nM completely prevented cell clumping and promoted the sprouting of neurite-like processes. Unexpectedly, cells treated with a single dose of 100 nM bufalin survived mostly intact for 4 weeks, whereas higher concentrations led to cell degeneration and death (Supp. Fig. 3B). Other cardiac glycosides, including oleandrin (100 nM), digoxin (100 nM), and proscillaridin A (3 nM), also prevented clumping, promoted neurite-like processes, and sustained differentiation over 4 weeks (Supp. Fig. 3C).

Next, we finalized a novel differentiation protocol involving two steps. SK-N-AS cells were plated at ∼10% confluence and allowed to proliferate in DMEM, 10% FBS to ∼30% confluence over 3 days. In stage 1 (S1), the medium was switched to DMEM, 1% FBS, and 5 µM PP121 for 4 days. In stage 2 (S2), the medium was switched to neurobasal, 1X B27, 0.25X Glutamax-I, 5 µM PP121, and 100 nM bufalin. New S2 medium (20% of the initial volume) was added weekly to replace the evaporated medium (Fig. 2A). To determine how long the differentiated cells could survive, we imaged them continuously every 4 h for 8 weeks (Fig. 2B, Supp. Video 1). Differentiated cells appeared mostly healthy for up to 4 weeks, with signs of degeneration slowly accumulating over the next 4 weeks. Intriguingly, over time, two distinct populations of cells emerged: a majority of non-motile, adherent, senescent-like cells and a minority of locally motile, poorly adherent, neuron-like cells that switched back and forth (Fig. 2B-C, Supp. Fig. 4, Supp. Video 1). Compared to naïve cells, immunostaining of S2D14 cells revealed a complete disappearance of the proliferation marker KI67 (Fig. 2C, D) and an increase in the length of neuronal marker TUJ1^+^ processes (Fig. 2C, E), suggesting possible neuronal differentiation.

**Figure 2:**
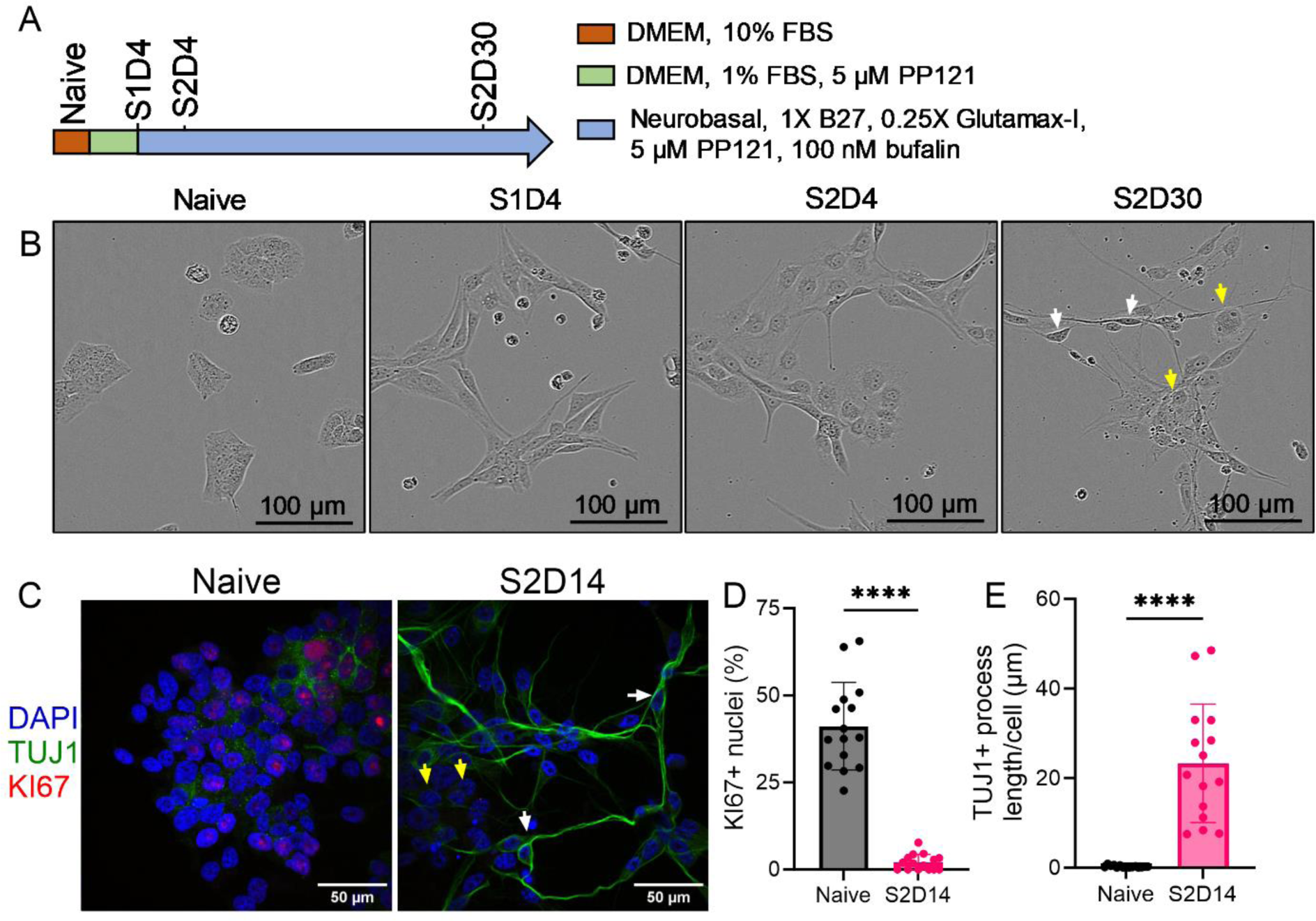
PP121 and bufalin synergistically differentiate SK-N-AS cells. **(A)** Schematic of optimized differentiation protocol for SK-N-AS cells. In stage 2 (S2), 20% of S2 medium is added every 7 days to replace the amount evaporated. **(B)** Representative images of cells over 30 days of differentiation. **(C)** Representative immunofluorescent images of naive and S2D14 cells. **(D-E)** Quantification of KI67-positive nuclei and TUJ1-positive processes (n=15 images per group, Student’s t-test). Error bars represent S.D.. The white arrows point to neuron-like cells with small cell bodies and neurite-like processes; the yellow arrows point to senescent-like cells with enlarged cell bodies and no processes. Source data are provided in the source data file.

### PP121/bufalin inhibits proliferation but activates a non-neuronal differentiation

To understand the molecular mechanisms contributing to PP121/bufalin-induced differentiation of SK-N-AS cells, we performed RNA-seq analyses on 4 groups of samples (N=4 each): naive, S1D4, S2D4, and S2D30. Compared to naïve cells, differentiated cells showed a progressive downregulation of specific genes involved in cell proliferation, such as DNA Topoisomerase II Alpha (TOP2A) and Marker Of Proliferation Ki-67 (MKI67) (Fig. 3A-C). This pattern was in line with the observed cell proliferation arrest. Interestingly, Paired Related Homeobox 1 (PRRX1), a transcription factor (TF) associated with MES cells (*7, 8, 35*), was the second most significantly upregulated gene between naïve and S1D4 cells (Fig. 3A). In contrast, Paired Like Homeobox 2A (PHOX2A) and Paired Like Homeobox 2B (PHOX2B), TFs associated with ADRN cells, were progressively downregulated in stage 2 (i.e., S2D4 and S2D30) (Fig. 3B-C). Unexpectedly, although PP121 promoted morphological changes that resembled neuronal differentiation, genes associated with neuronal states, such as microtubule-associated protein 1A (MAP1A) and synaptophysin (SYP), were downregulated in S2D4 cells (Fig. 3B). Genes associated with inflammation, such as C-C Motif Chemokine Ligand 3 (CCL3) and C-C Motif Chemokine Ligand 3 Like (3CCL3L3), were upregulated in S2D30 cells (Fig. 3C).

**Figure 3:**
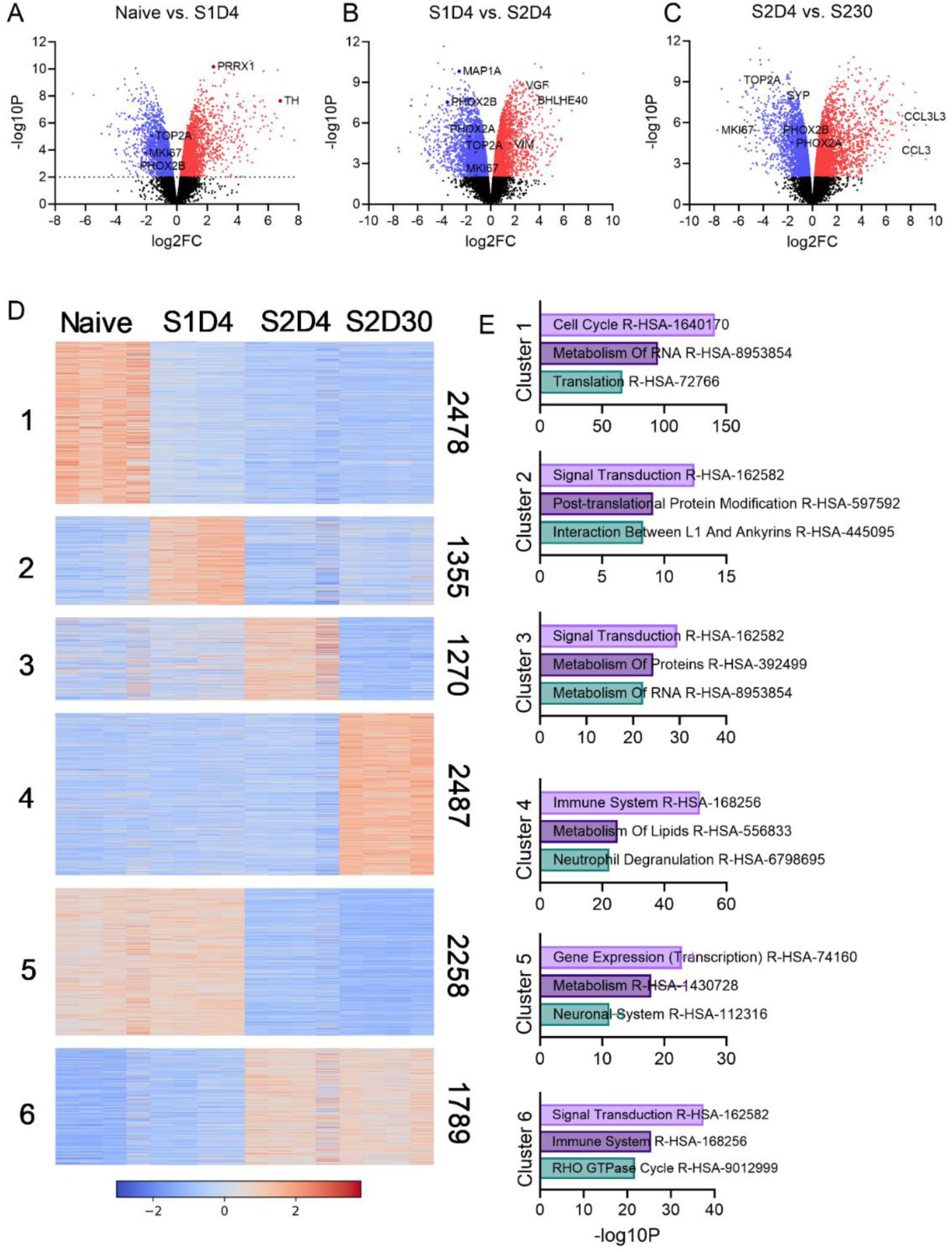
PP121/bufalin drives transcriptional changes associated with non-neuronal differentiation of SK-N-AS cells. (A-C) Volcano plots comparing different stages of SK-N-AS differentiation. Significantly downregulated genes are colored blue, and significantly upregulated genes are colored red (N=4 per stage). (D) Heatmap of all differentially expressed genes across four states. Z-score scaling was applied internally for each gene before K-means clustering. Numbers on the left indicate the cluster number, and numbers on the right indicate the number of genes in that cluster (refer to Table S3 for the complete list of genes in each cluster). (E) Top Reactome pathway enrichment analyses for each cluster in D.

Next, to understand the trajectory of differential gene expression, we performed K-means clustering of differentially expressed genes (P <0.01) that resulted in 6 distinct clusters (Fig. 3D, Supp. Table 3), and analyzed each cluster for Reactome pathway enrichment (Fig. 3E) using Enrichr (*36*). Cluster 1, which was progressively downregulated over time, was enriched for pathways associated with Cell Cycle, Metabolism of RNA, and Translation, supporting the observation of complete arrest of cell proliferation. Cluster 2 (Fig. 3F), containing genes upregulated at S1D4, was enriched for the Interaction Between L1 and Ankyrins, which is associated with neurite outgrowth (*37*). Cluster 3, containing genes upregulated at S2D4, was enriched for Metabolism of RNA and Proteins. Cluster 4, containing genes upregulated at S2D30, was enriched for Immune System and Neutrophil Degranulation, suggesting an increased inflammatory state. Cluster 5, for genes downregulated in both S2D4 and S2D30, was enriched for Transcription, Metabolism, and, unexpectedly, Neuronal System. Cluster 6, for genes upregulated in both S2D4 and S2D30, was enriched for Immune System and RHO GTPase Cycle, which regulates many pathways, including cytoskeleton reorganization and cell adhesion (*38*). Interestingly, Signal Transduction was enriched in clusters 2, 3, and 6, suggesting increased extracellular communications across differentiation states. Together, transcriptomic analyses supported a global reduction in cell division, transcription, and translation, and a global increase in extracellular communication and inflammation. However, contrary to their neuron-like morphology, long-term differentiated cells did not appear to transcriptionally acquire a neuronal fate (cluster 5).

### PP121/bufalin leads to a collapse of ADRN CRC and a gain of MES CRC

Considering the upregulation of MES-associated TF PRRX1 in S1D4 and the downregulation of ADRN-associated PHOX2B in S2D4 and S2D30, we visualized additional previously identified TFs associated with MES or ADRN transcriptional programs (*7*). A majority of ADRN TFs appeared to be steadily downregulated over time (Fig. 4A). Interestingly, MES TFs appeared to be upregulated in waves, with a specific subgroup of genes upregulated at a particular stage (Fig. 4B). Next, we combined our data with two published neuroblastoma datasets (N=67 cell lines) (*6, 39*) and performed gene signature analysis as previously described (*7*). As expected, MES cell lines had high MES scores and low ADRN scores and closely resembled neural crest cells, whereas ADRN cell lines had low MES scores and high ADRN scores (Fig. 4C). Naive SK-N-AS cells exhibited intermediate ADRN-MES state (*6*) and became progressively more MES-like and less ADRN-like at differentiation (Fig. 4C). As a recent study found that MES-type cells were more immunogenic than ADRN-type cells (*40*), we performed similar analyses for immune activation and immune evasion gene sets. As expected, differentiated SK-N-AS cells also became more enriched for immune activation and evasion genes set (Supp. Fig. 5A). Furthermore, as cells were in culture for more than 30 days, we hypothesized that they also became senescent. Using a recently established senescent gene set (*41*), we found that differentiated SK-N-AS cells indeed became significantly more senescent and also found a novel positive correlation between MES and senescence scores (Supp. Fig. 5B).

**Figure 4:**
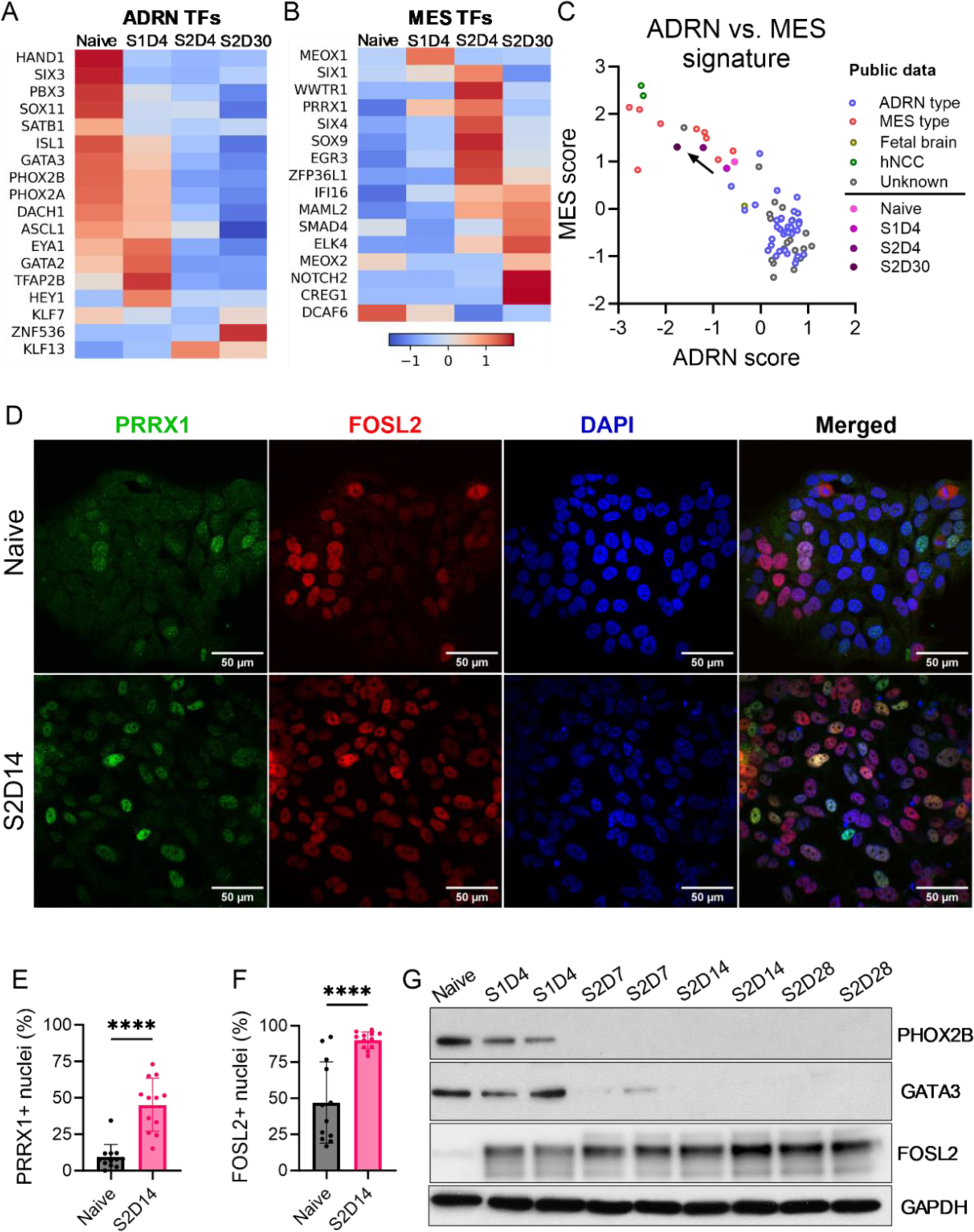
PP121/bufalin leads to a collapse of ADRN CRC and a gain of MES CRC. **(A-B)** Averaged heatmap (N=4) for ADRN and MES TFs. Z-score scaling was applied internally for each gene. **(C)** Differentiated SK-N-AS cells became increasingly more MES and less ADRN (From public databases, N =67 samples; from this study, N=4 per group). **(D)** Representative IF images for naïve and S2D14 SK-N-AS cells. **(E-F)** Quantification of PRRX1-positive and FOSL2-positive nuclei (N=12 images per group, Student’s t-test). **(G)** Representative Western blots of naïve and differentiated SK-N-AS cells over time. Error bars represent S.D. Source data and uncropped Western blots are provided in the source data file.

To validate RNA-seq analyses at the protein level, we performed immunofluorescent imaging (IF) for naïve and S2D14 cells with antibodies for MES-associated TFs PRRX1 and FOS like 2, AP-1 transcription factor subunit (FOSL2) (Fig. 4D). As an ADRN-MES intermediate cell line, 9.2% and 47% of naïve SK-N-AS cell nuclei were positive for PRRX1 and FOSL2, respectively, which increased to 45% and 90% in S2D14 cells (Fig. 4E-F). Western blots of naïve and differentiated SK-N-AS cells also showed a complete loss of ADRN-associated TFs PHOX2B and GATA-binding protein 3 (GATA3) starting at S2D7 and a steady increase in FOSL2 starting at S1D4 (Fig. 4G). Therefore, differentiated SK-N-AS cells also showed a gain in MES-associated TFs and a loss of ADRN-associated TFs at the protein levels, which is in agreement with our RNA-seq data.

### PP121/bufalin upregulates transcription factors associated with SCPs and chromaffin cells

Various studies have established core regulatory circuits (CRCs) that define neuroblastoma identity, including a retino-sympathetic CRC associated with RA-induced differentiation (*28*). To determine if our PP121/bufalin-treated cells resembled those differentiated by RA, we compared Reactome pathways represented by down- or upregulated genes in our cells (S2D30 vs. naive) with corresponding datasets in neuroblastoma cell lines differentiated by RA (*28*). There was a significant overlap in pathways of genes downregulated by RA and by PP121/bufalin-differentiated SK-N-AS cells, primarily pathways associated with cell division (Fig. 5A, Supp. Fig. 6A). However, there was much less overlap in the pathways of upregulated genes (Fig. 5B, Supp. Fig. 6B). TFs associated with the retino-sympathetic CRC previously established (*28*) were also downregulated or unchanged, further supporting that PP121/bufalin-induced differentiation was distinct from RA-induced differentiation (Fig. 5C). A previous study of the developing human adrenal glands (*29*) showed that cycling cells could differentiate into late neuroblasts, chromaffin cells, or Schwann cell precursors (SCPs). As RA was shown to differentiate neuroblastoma cells towards neuroblasts, we hypothesized that our differentiated cells might instead resemble chromaffin cells or SCPs. Indeed, TFs driving developmental programs in chromaffin cells (*29*) were significantly upregulated by our differentiation protocol (Fig. 5D).

**Figure 5:**
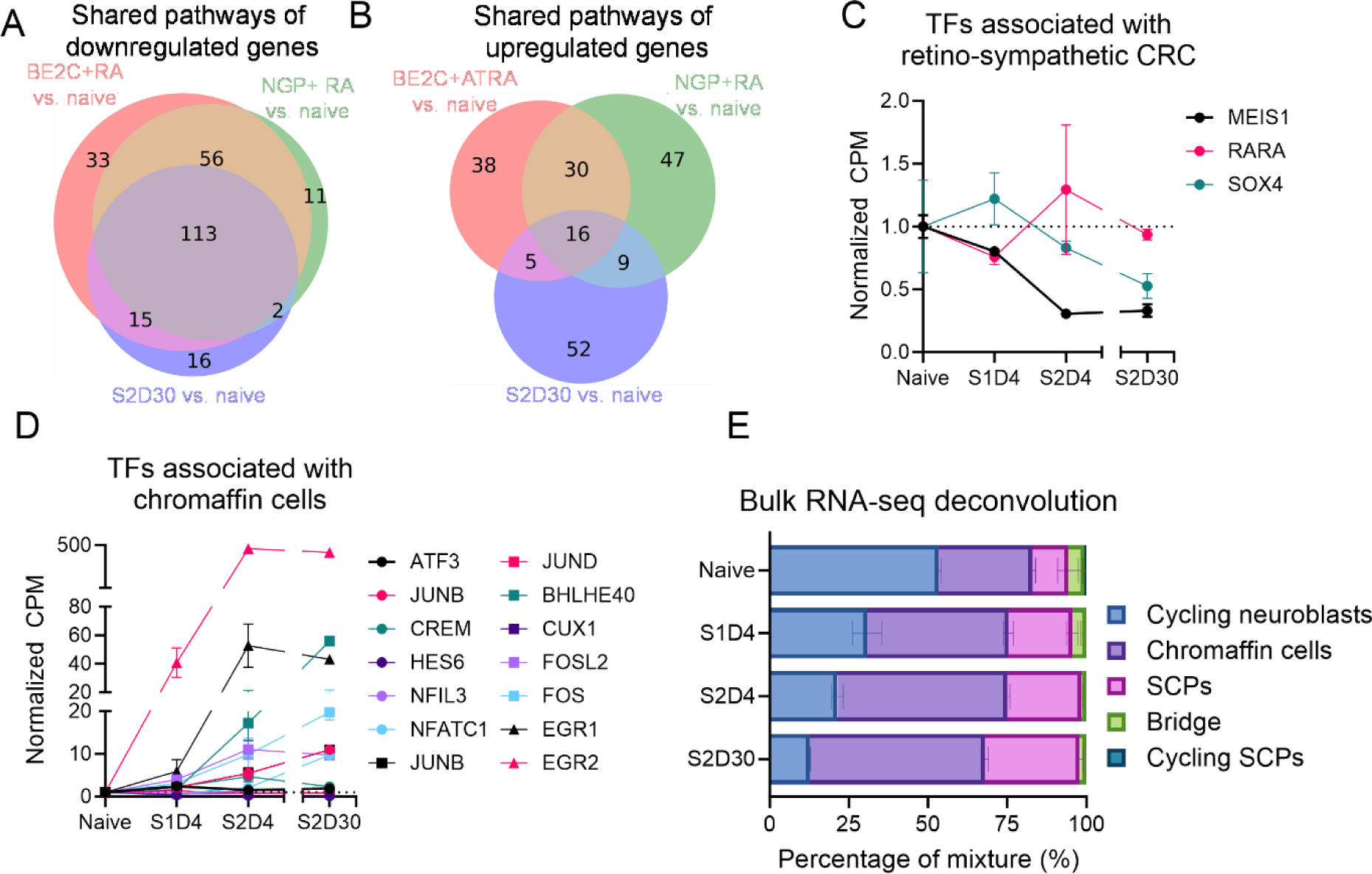
Differentiated SK-N-AS cells show a gain in TFs associated with chromaffin and SCP state. **(A-B)** Shared down- and upregulated pathways among SK-N-AS cells differentiated with PP121/bufalin, and BE2C and NGP cells differentiated with RA. **(C)** TFs associated with retino-sympathetic CRC were downregulated or unchanged. **(D)** TFs associated with chromaffin cells were generally upregulated. **(E)** Deconvolution of bulk RNA-seq results of naïve and differentiated SK-N-AS cells (N=4). Error bars represent S.D.. Source data are provided in the source data file.

Next, we utilized an unbiased approach to determine the developmental states imposed by PP121/ bufalin. We used Cibersort (*42*) to generate a reference matrix atlas from the single-cell RNA-seq data in the human sympathoadrenal development study (*29*) (Supp. Table 4). With this atlas, we then deconvoluted our bulk RNA-seq data to infer developmental cell states. Naïve SK-N-AS cells were predicted to contain a mix of cycling neuroblasts, chromaffin cells, SCPs, and bridge cells. Over the course of differentiation, the percentage of cycling neuroblasts decreased, whereas the percentage of chromaffin cells and SCPs increased (Fig. 5E). However, at the RNA level, while some markers of mature chromaffin cells and SCPs increased in S1D4 and S2D4, the majority decreased in S2D30 (Supp. Fig. 6C, D), suggesting that these cells resembled intermediate forms of chromaffin cells and SCPs.

### PP121/bufalin promotes long-term differentiation across multiple cell lines

We further observed that the switch to neurobasal was not necessary for differentiation. Instead, SK-N-AS cells can be differentiated in one step by adding 5 µM PP121 and 10 nM bufalin directly to the growth medium (Fig. 6A, Supp. Video 2). Applying this one-step protocol, we successfully induced long-term differentiation of additional MES-type neuroblastoma GI-ME-N and SH-EP cells with 2.5 µM PP121 and 10 nM bufalin (Fig. 6B, Supp. Fig. 7A-C, Supp. Videos 3-4) for up to 16 weeks. Using a similar protocol, we also differentiated and maintained glioblastoma U-251 and LN-229 cells with 5 µM PP121 and 30 nM bufalin (Fig. 6B, Supp. Videos 5-6) for 16 weeks. As previously observed, the addition of PP121 killed ADRN-type SK-N-SH, Kelly, and SH-SY-5Y cells (Supp. Fig. 7D-F, Supp. Videos 7-9). While the cells did not clump extensively without bufalin in DMEM or RPMI medium, bufalin further-improved long-term health for GI-ME-N and SH-EP cells. Interestingly, bufalin was necessary to completely inhibit the proliferation in LN-229 and U-251 cells (Supp. Fig. 7G-H, Supp. Videos 5-6). Day 14 (D14) differentiated GI-ME-N cells showed a loss of MKI67 staining, narrow cell bodies, and a more elaborated network of TUJ1^+^neurite-like processes (Fig. 6C, E). D14 differentiated SH-EP cells also showed a loss of MKI67, more elongated shapes, and increased TUJ1 staining. However, no distinct processes were observed (Fig. 6D, F). Similar to SK-N-AS cells, deconvolution of bulk RNA-seq data from naive and D30 GI-ME-N cells showed a loss of cycling neuroblasts and cycling SCPs and a gain of chromaffin cells and SCPs (Fig. 6G).

**Figure 6:**
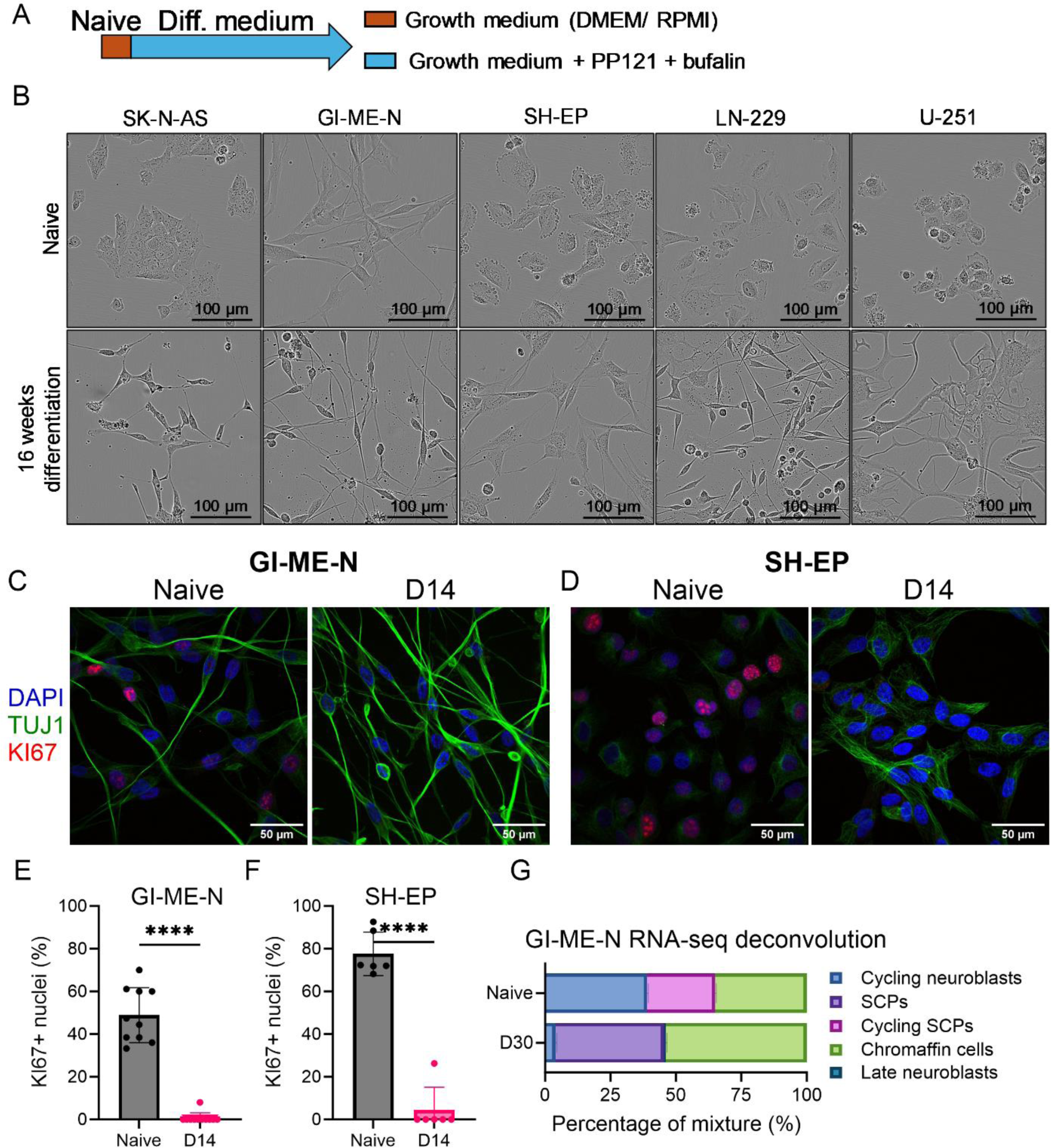
PP121/bufalin promotes the differentiation of multiple cell lines. **(A)** Schematic of the one-step differentiation protocol. **(B)** Representative live cell images of SK-N-AS, GI-ME-N, SH-EP, LN-229, and U-251 cells after 16 weeks of differentiation. **(C-D)** Representative IF images of naïve and D14 GI-ME-N and SH-EP cells. **(E-F)** Quantification of KI67-positive nuclei in GI-ME-N (N=12) and SH-EP (N=6) cells. **(G)** Deconvolution of bulk RNA-seq results of naïve and D30 differentiated GI-ME-N cells (N=3). Error bars represent S.D.. Source data are provided in the source data file.

### Possible mechanisms of PP121/bufalin-induced differentiation

Although the observed morphological changes are striking and consistent across cell lines, the underlying mechanisms remain unknown. We conceptualized three potential complementary mechanisms driving cell differentiation. First, we hypothesize that PP121 induces differentiation by upregulating miR-124. miR-124 was indeed consistently upregulated in differentiated SK-N-AS and GI-ME-N cells (Fig. 7A, C). Notable direct targets of miR-124 that were downregulated with differentiation include cyclin-dependent kinases 4 and 6 (CDK4, CDK6) (*43, 44*). Other downregulated miR-124 targets include Enhancer of zeste homolog 2 (EZH2) (*45*) and PHD finger protein 19 (PHF19) (*46*), which are components of the Polycomb Repressive Complex 2 (PRC2) (Fig. 7A, C) that regulate histone methylation and transcriptional repression. These observations suggest that miR-124 plays a role in inhibiting proliferation and driving differentiation.

**Figure 7:**
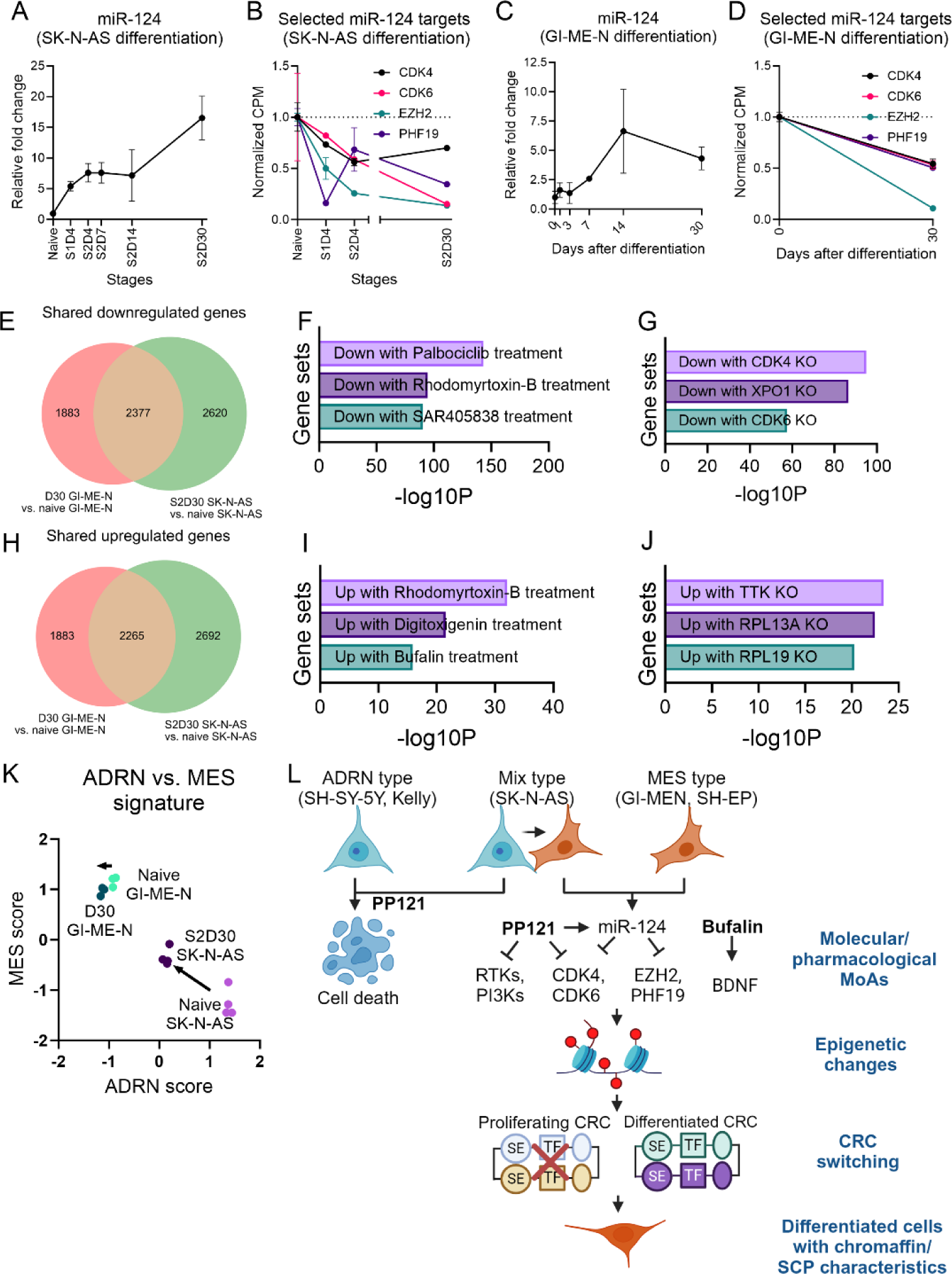
Proposed possible mechanisms for PP121/bufalin-induced differentiation of MES-type neuroblastoma cells. **(A-B)** miR-124 was upregulated and some of its targets were downregulated in SK-N-AS cells. **(C-D)** Similar observations for GI-MEN cells. **(E-K)** Consensus molecular drug and gene CRISPR KO signatures for shared downregulated genes. **(H-J)** Consensus molecular drug and gene CRISPR KO signatures for shared upregulated genes. **(K)** ADRN and MES signature score for naïve and differentiated SK-N-AS and GI-ME-N cells. **(L)** Proposed mechanism for PP121/bufalin-induced differentiation of MES-type neuroblastoma cells. Error bars represent S.D.. Source data are provided in the source data file. Fig. 7L was created with BioRender.com.

Second, PP121, a dual inhibitor targeting tyrosine kinases and phosphoinositide kinases (*27*), and bufalin, a Na^+^/K^+^ ATPase inhibitor (*47*), may induce differentiation through their established and miRNA-independent mechanisms of action. To test this hypothesis, we first determined the shared significantly down- and upregulated genes (P< 0.01) in terminally differentiated SK-N-AS cells (S2D30) and GI-ME-N cells (D30) relative to their naïve counterparts, and then used Enrichr (*36*) to determine enrichment signatures for drugs and gene-specific CRISPR knockout (KO) (Fig. 7E-L). Interestingly, shared downregulated genes were significantly enriched for genes downregulated by Palbociclib (Fig. 7F), a CDK4/6 inhibitor recently shown to promote the differentiation of ADRN cells but not MES cells (*48*). Shared downregulated genes were also enriched for genes downregulated by CDK4/6 CRISPR KO (Fig. 7G). Shared upregulated genes were enriched for genes upregulated by cardiac glycosides digoxigenin and bufalin (Fig. 7I) and by CRISPR KO of dual-specific protein kinase TTK (TTK) (Fig. 7J). Notably, shared down- and upregulated genes were also enriched for genes down- and upregulated by Rhodomyrtoxin-B, an antibacterial and cytotoxic compound predicted to be a DNA intercalator (*49*). These observations support that the known mechanisms of PP121 and bufalin drive parts of the changes in gene expressions observed, as well as further emphasizing CDK4/6 inhibition as an important part of the mechanism.

Third, epigenetic modifications, both miR-124-dependent and independent, likely play a significant role. Differentiation of ADRN-type cells by RA involved epigenetic reprogramming (*28*), and miR-124 is known to drive neuronal differentiation through epigenetic mechanisms (*50*). We also observed nuclei elongation in differentiated cells as measured by increased nuclei eccentricity (Supp. Fig. 8A-H), which may be a response to differentiated cells also becoming more elongated. Both elongated cell and nuclei morphology have been associated with differentiation and chromatin remodeling in mesenchymal (*51*) and epidermal (*52*) stem cells, further supporting that epigenetic modifications play an essential role in PP121/bufalin-induced differentiation.

Besides potential mechanisms, we observed that PP121 killed ADRN-type cells while promoting differentiation in mixed SK-N-AS cells and MES GI-MEN and SH-EP cells. Differentiated SK-N-AS cells exhibited a large shift towards increased MES and decreased ADRN characteristics (Fig. 4, Fig. 7K). However, differentiated GI-ME-N cells did not show a significant change in MES or ADRN scores compared to their naïve counterparts (Fig. 7K). These findings suggest that PP121/bufalin kills ADRN-type cells or forces their conversion to MES-type cells, while MES-type cells survive and differentiate, resulting in the observed phenotypic shift in SK-N-AS cells, but not in GI-ME-N cells.

We propose a synthesis of these pathways, as illustrated in Fig. 7L. PP121/bufalin specifically induces the differentiation of MES-type cells through their established mechanisms of action as well as through miR-124 upregulation. This results in epigenetic changes that lead to permanent alterations of CRCs, switching them from proliferative to differentiated states. The final outcome is the differentiation of MES neuroblastoma cells into non-proliferating cells with chromaffin and SCP characteristics. The specific components of this scheme remain to be tested and refined.

## DISCUSSION

Here, we presented a novel strategy for inducing and maintaining the differentiation of SK-N-AS cells over prolonged periods by harnessing the synergistic effects of two compounds - PP121 and bufalin. PP121, a multikinase inhibitor, functioned to arrest cell proliferation and facilitated the sprouting of cellular processes, while bufalin, a cardiac glycoside, prevented cell clumping and promoted sustained cell health. PP121 and bufalin also differentiated other MES neuroblastoma cells, including GI-ME-N and SH-EP cells, and glioblastoma cells, including U-251 and LN-229 cells. The drug combination not only triggered cellular differentiation but also preserved the specialized phenotype for an extended duration, with cells maintaining their distinctive characteristics for at least 16 weeks. Notably, various evidence suggests that PP121/bufalin-differentiated MES neuroblastoma cells resembled chromaffin cells and SCPs, which were distinct from the late neuroblast subtype promoted by RA (*29*). The simplicity and versatility of this method present a promising pathway for future research and potential clinical application for high-risk, MES-type neuroblastoma that exhibits resistance to conventional treatment modalities. The long-lived cells generated can also serve as valuable models for studying various processes such as differentiation, maturation, inflammation, senescence, and response to therapeutic agents.

While further investigation is required to explore the proposed pathways underlying PP121/ bufalin-induced differentiation (Fig. 7L), unraveling these mechanisms will provide valuable insights into neuroblastoma cell fate decisions and new therapeutic strategies. During the development of the adrenal gland, there is a gradual loss of cycling neuroblasts, which, when abberantly persisted, are associated with high-risk neuroblastoma (*29*). PP121/bufalin-induced differentiation appears to mimic this process by reducing the population of cycling neuroblasts (Fig. 5E) and cycling SCPs (Fig. 6G), and therefore potentially lowering disease aggressiveness. Recent genetic studies suggest that SCPs are generated during early development and can give rise to chromaffin cells and neuroblasts (*29, 53*). In the SK-N-AS cell model, we further observe a decrease in bridge cells (*29*), which are intermediates connecting SCPs to chromaffin cells, with differentiation. These observations suggest that PP121/bufalin-induced differentiation partially resembles normal cellular differentiation in fetal adrenal gland development. It should be noted that chromaffin cells in the adrenal medulla lack dendrites or axons (*5*). Nevertheless, our differentiated cells exhibit both enriched chromaffin/SCP genetic signatures and enhanced neuronal morphology, such as TUJ1-positive processes. Therefore, these differentiated cells have features related to multiple developmental states. Whether this complex differentiation pattern is an artifact of the drug combination treatment or resembles normal development remains to be investigated.

One significant limitation of our study is the inability of bulk RNA-seq derived from pooled cell populations (*54*) to fully capture the heterogeneity of the neuroblastoma cell population, particularly evident in SK-N-AS cells, which comprise a mixture of ADRN and MES-type cells. Notably, time-lapse videos revealed diverse cellular morphologies in differentiated SK-N-AS cells, while differentiated GI-ME-N and SH-EP cells, being predominantly MES-type, appeared more homogeneous. Overcoming these limitations through single-cell analysis is important for comprehending the dynamic processes underlying neuroblastoma progression and differentiation. Future directions also entail a thorough exploration of the mechanisms driving cell differentiation (Fig. 7L), particularly epigenetic mechanisms. Furthermore, several studies have supported that cardiac glycosides, including bufalin, are potent anti-cancer compounds (*47, 55*) but are toxic at high doses (*56*). While *in vivo* data for the efficacy of PP121 is relatively sparse, it was shown to reduce tumor growth in prostate (*57*), thyroid (*58*), and esophageal (*59*) in mouse cancer models. To our knowledge, a combination of PP121/bufalin has not been tested in animal models, which will be necessary to validate observed effects and contextualize therapeutic potential. By leveraging mechanistic insights and preclinical testing, PP121/bufalin-induced differentiation can be potentially developed to inhibit the growth of neural cancers while minimizing toxicity, especially in pediatric patients (*60*), thereby mitigating long-term side effects associated with conventional treatments (*61*).

In conclusion, our innovative protocol offers a promising avenue for treating therapy-resistant neuroblastoma. The long-lived, differentiated cells can be valuable tools for investigating various cellular processes. Therefore, our findings contribute to advancing both basic scientific knowledge and clinical interventions.

## Supporting information

Differentiation of SK-N-AS cells over 56 days

SK-N-AS cells treated with vehicle (0.1% DMSO) over 7 days

SK-N-AS cells treated with 5 uM PP121 over 7 days

SK-N-AS cells treated with 5 uM PP121 + 10 nM bufalin over 7 days

GI-ME-N cells treated with vehicle (0.1% DMSO) over 7 days

GI-ME-N cells treated with 2.5 uM PP121 + 10 nM bufalin over 7 days

GI-ME-N cells treated with 2.5 uM PP121 over 7 days

SH-EP cells treated with vehicle (0.1% DMSO) over 7 days

SH-EP cells treated with 2.5 uM PP121 over 7 days

SH-EP cells treated with 2.5 uM PP121 + 10 nM bufalin over 7 days

LN-229 cells treated with vehicle (0.1% DMSO) over 7 days

LN-229 cells treated with 5 uM PP121 over 7 days

LN-229 cells treated with 5 uM PP121 + 30 nM bufalin over 7 days

U-251 cells treated with vehicle (0.1% DMSO) over 7 days

U-251 cells treated with 5 uM PP121 over 7 days

U-251 cells treated with 5 uM PP121 + 30 nM bufalin over 7 days

Kelly cells treated with vehicle (0.1% DMSO) over 7 days

Kelly cells treated with 5 uM PP121 over 7 days

Kelly cells treated with 5 uM PP121+ 10 nM bufalin over 7 days

SH-SY-5Y cells treated with vehicle (0.1% DMSO) over 7 days

SH-SY-5Y cells treated with 5 uM PP121 over 7 days

SH-SY-5Y cells treated with 5 uM PP121+10 nM bufalin over 7 days

SK-N-SH cells treated with vehicle (0.1% DMSO) over 7 days

SK-N-SH cells treated with 5 uM PP121 over 7 days

SK-N-SH cells treated with 5 uM PP121+ 10 nM bufalin over 7 days

## Acknowledgments

The NeuroTechnology Studio at Brigham and Women’s Hospital provided IncuCyte instrument access and consultation on data acquisition and analysis. Biorender was used in the preparation of Figure 7.

## Funding sources

AMK was funded by the Rainwater Charitable Foundation and NIH NCI 1R01-CA215072.

REG was funded by the NIH R01-CA271605.

SS was funded by a research grant from the Friends of Dana-Farber Cancer Institute.

## Author contributions

LDN, SS, and AMK conceived and designed the study.

LDN, SS, KC, and AF performed experiments and analysis for this study.

REG and AMK provided the resources needed for experiments.

LDN wrote an original draft of the manuscript.

LDN, SS, KC, and AMK reviewed and edited the manuscript.

All authors reviewed and commented on the manuscript.

## Potential conflict of interests

The authors have no conflicts of interest to declare.

## Data and Material Availability

mRNA-sequencing data that support the findings of this study were deposited into the Gene Expression Omnibus (GEO) Repository with accession number GSE270493. Relevant CPM values for all analyses performed are provided in Supp. Table 8. Source data for all relevant figures are provided in the source data file. Contact the corresponding author to request materials and cell lines used in the manuscript.

## MATERIALS AND METHODS

### Cell Cultures

SK-N-AS, SK-N-SH, SH-SY-5Y, Kelly, LN-229, and U-251 cells were maintained in DMEM+ GlutaMAX™ supplemented with 10% fetal bovine serum (FBS) and 1% penicillin/streptomycin (P/S). GIMEN and SH-EP were maintained in RPMI 1640 Medium + GlutaMAX™ supplemented with 10% FBS and 1% penicillin/streptomycin (P/S). Cells were cultured in a standard incubator set to a controlled environment of 5% CO2 and 37°C and checked for mycoplasma contamination every 3 months.

The protocol for primary rat neuron preparation was approved by the Institutional Animal Care and Use Committee at Brigham and Women’s Hospital. Rat primary cortical neuron cultures were prepared from E18 SAS Sprague Dawley pups (Charles River). Brain tissues were dissected, dissociated enzymatically by 0.25% Trypsin-EDTA (Thermo Fisher Scientific), triturated with fire-polished glass Pasteur pipettes, and passed through a 40 μm cell strainer (Sigma-Aldrich). After counting, neurons were seeded onto poly-D-lysine (Sigma-Aldrich) coated cell culture plates at 80,000 cells/cm^2^ in Neurobasal medium supplemented with 1X B27 and 0.25X GlutaMax. Half medium was changed every 4 days until use.

### miR-124 Transfection

miR-124 mimics or non-targeted CTRL (Horizon Discovery) were dissolved in water to 50 µM concentration and used at the final concentration of 50 nM. Transfection into cultured SK-N-AS cells was performed using Lipofectamine 2000 (Thermo Fisher Scientific) following standard protocols. After a 24-hour incubation period, the transfection medium was replaced with a fresh growth medium, and cells were subsequently analyzed for confluence and viability 4 or 7 days after transfection.

### Incucyte Live Imaging

Live cell imaging was performed using the IncuCyte^TM^ Live-Cell Imaging System (Sartorius) at 20X magnification. Cell confluence was quantified using the IncuCyte^TM^ software.

### WST-1 Cell Viability Assay

Cell viability was measured by WST-1 reduction assay (Sigma-Aldrich). For the assay, all medium was removed and replaced with 1X WST-1 reagent dissolved in the growth medium, followed by 1 hour of incubation at 37°C. The absorbance of the culture medium was measured with a microplate reader at test and reference wavelengths of 450 nm and 630 nm, respectively. Relative viability values were obtained by normalizing to the CTRL mimics or vehicle conditions.

### Drug Treatment

PP121 (Medchemexpress) was dissolved to 10 mM in DMSO and used at final concentrations of 2.5 – 10 µM. Bufalin (Fisher Scientific) was dissolved to 1 mM in DMSO and used at final concentrations of 10 – 100 nM. Detailed concentrations used to differentiate each cell line were listed in Supp. Table 5.

### RNA Collection and RT-qPCR

Total RNA from cells was extracted using the Norgen Total RNA Purification Kit (Norgen Biotek) following the manufacturer’s protocol. DNAse1 was applied during RNA extraction to remove genomic DNA. RNA was eluted in nuclease-free water, and the concentration was measured using Nanodrop (Thermo Fisher Scientific). For miRNA analysis, 50ng of RNA was reverse-transcribed into cDNA using the miRCURY LNA RT kit (Qiagen). RT-qPCR mix was prepared using the miRCURY LNA SYBR Green PCR kit (Qiagen). qPCR was performed using the QuantStudio 7 Flex System. The cycling conditions were 95°C for 10 min, 50 cycles of 95°C for 15 s, and 60°C for 1 min following dissociation analysis. miRNA expression was normalized to let-7a. Quantification was performed using the delta-delta Ct method. miRNA and mRNA primers used were listed in Supp. Table 6.

### Immunofluorescent Staining and Analysis

Cells plated on coverslips were fixed with 4% formaldehyde in PBS (Corning) for 15 min, washed in PBS, incubated in 1X blocking/permeabilization buffer (Biotium) for 1h, and incubated with primary antibodies overnight at 1:100 dilution. Coverslips were washed with PBS and incubated with the corresponding AlexaFluor-conjugated secondary antibodies at 1:500 dilution (Life Technologies) for 2h at room temperature. After washing, coverslips were mounted onto glass slides containing DAPI mounting medium. Image acquisition was done with a Zeiss LSM 710 Confocal Microscope. Primary and secondary antibodies used were listed in Supp. Tables 7. MKI67 expression and morphology analyses were conducted with CellProfiler software (*62*). Custom pipelines were developed for each cell line, with the same pipeline used for naïve and differentiated cells of the same line.

### RNA-seq and Transcriptomic analyses

After quality control by Agilent 2100 Bioanalyzer, the total RNA was used as input for library preparation by Novogene Co., Ltd, followed by high-throughput sequencing on Illumina HiSeq X with PE150 mode to produce approximately 20 M reads per sample. Raw data alignment and differential gene expression analysis were performed using the BioJupies web application (*63*). For heatmap and k-means clustering, the CPM values were first normalized internally for each gene. Python’s scikit-learn library was used to perform k-mean clustering into 6 clusters. Gene ontology analyses were performed using Enrichr web application (*36*).

### Gene Signature Analysis

Gene signature analysis utilized gene lists identified in previous studies: MES-specific and ADRN-specific genes (*7*), immune activation and evasion genes (*40*), and senescence genes (*41*). Genes were rank-ordered according to their CPM level, and the percentile was calculated relative to the full list. For each gene set, the percentile averages were Z-transformed across samples and then used as a signature score.

### Western Blots

Total protein was extracted using RIPA buffer (Boston Bioproducts) supplemented with Complete, Mini, EDTA-free Protease Inhibitor Cocktail (Millipore Sigma). Protein concentrations were determined using the Micro BCA Protein Assay Kit (Thermo Fisher Scientific). Equal amounts of protein were loaded, and electrophoresis was performed in NuPAGE 4 to 12% gradient Bis-Tris polyacrylamide protein gels (Thermo Fisher Scientific). Proteins were transferred to Immun-Blot PVDF membranes (Bio-Rad) and then blocked with 5% milk in tris-buffered saline with 0.1% Tween (TBS-T, Boston Bioproduct) for 1 h. Membranes were incubated overnight with primary antibodies at 4°C. Blots were washed and incubated with secondary antibodies for 2 h at room temperature. After washing, bands were visualized with ECL chemiluminescence reagents (Genesee Scientific) using the iBright Imaging System (Thermo Fisher Scientific). Band intensity was measured using the Image Studio Lite software (LI-COR Biosciences). Antibodies and other key resources are listed in Supp. Tables 7.

### Deconvolution of Bulk RNA-seq

Single-cell RNA-seq data from the developing human adrenal gland was downloaded (*29*) and used to prepare the reference matrix with the CIBERSORTx web application (*42*). This reference matrix was then used to deconvolute bulk RNA-seq data with CIBERSORTx web application.

### Statistical Analysis

Data management and calculations were performed using Prism 10 (GraphPad). Comparisons between the two groups were made using the unpaired two-tailed Student’s t-test. A P value < 0.05 was considered statistically significant, and the following notations are used in all figures: *P < 0.05, **P < 0.01, ***P < 0.001, and ****P < 0.0001. All error bars shown represent mean +/− standard deviation (SD) unless otherwise stated.

## Supplementary Figures

**Supplementary Figure 1:**
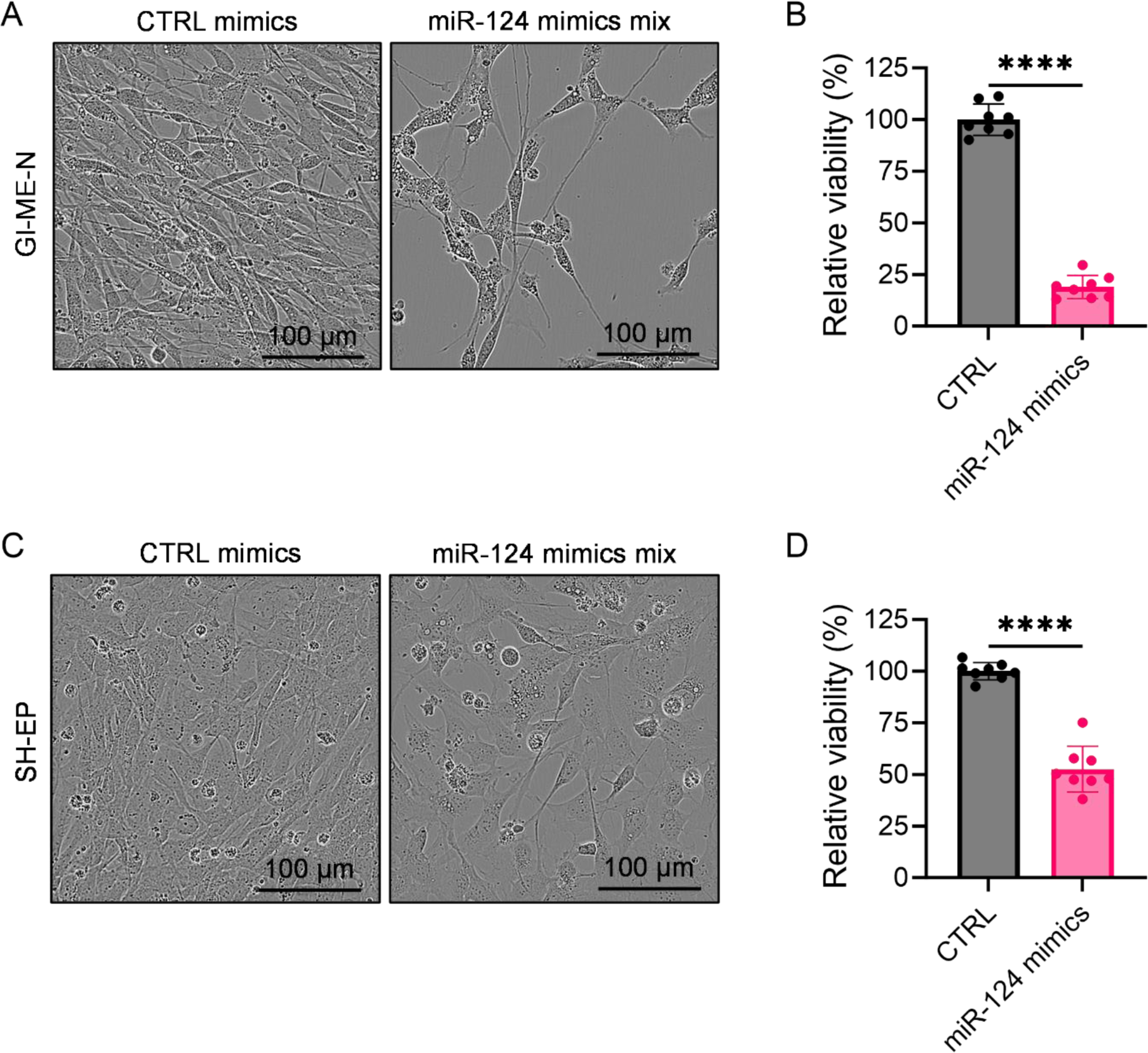
miR-124 transfection reduces proliferation in GI-ME-N and SH-EP cells. Transfection with miR-124 mimics (16.7 nM of each of the 3 pre-miR-124s) for 7 days reduced cell viability in GI-ME-N cells **(A-B)** and SH-EP cells **(C-D)** compared to the non-targeted CTRL (50 nM) (N=8 each, Student’s t-test). Error bars represent S.D.. Source data are provided in the source data file.

**Supplementary Figure 2:**
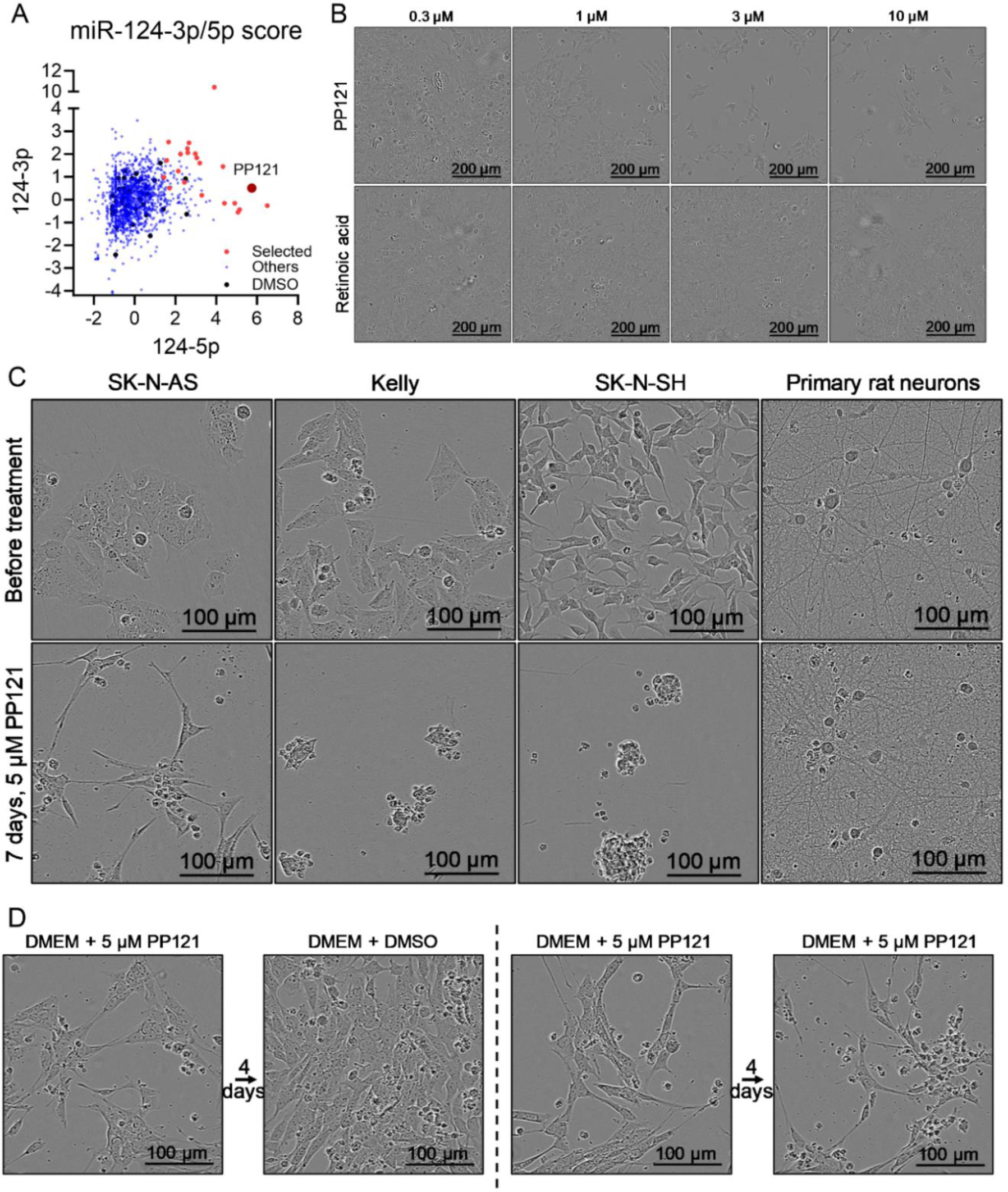
PP121 reversibly differentiates SK-N-AS cells. **(A)** Graph showing COMBAT score of all compounds, candidate compounds are shown in red, DMSO controls in black, and other compounds in blue. **(B)** Representative images of SK-N-AS cells 4 days after treatment with various doses of PP121 and retinoic acid (RA). **(C)** Representative images of different cell lines 7 days after treatment with 5 µM PP121. **(D)** Representative images of SK-N-AS cells 4 days after treatment with 5 µM PP121, followed by 4 days in fresh medium containing either DMSO or 5 µM PP121. Source data for 1A are included in Supplementary Table 1.

**Supplementary Figure 3:**
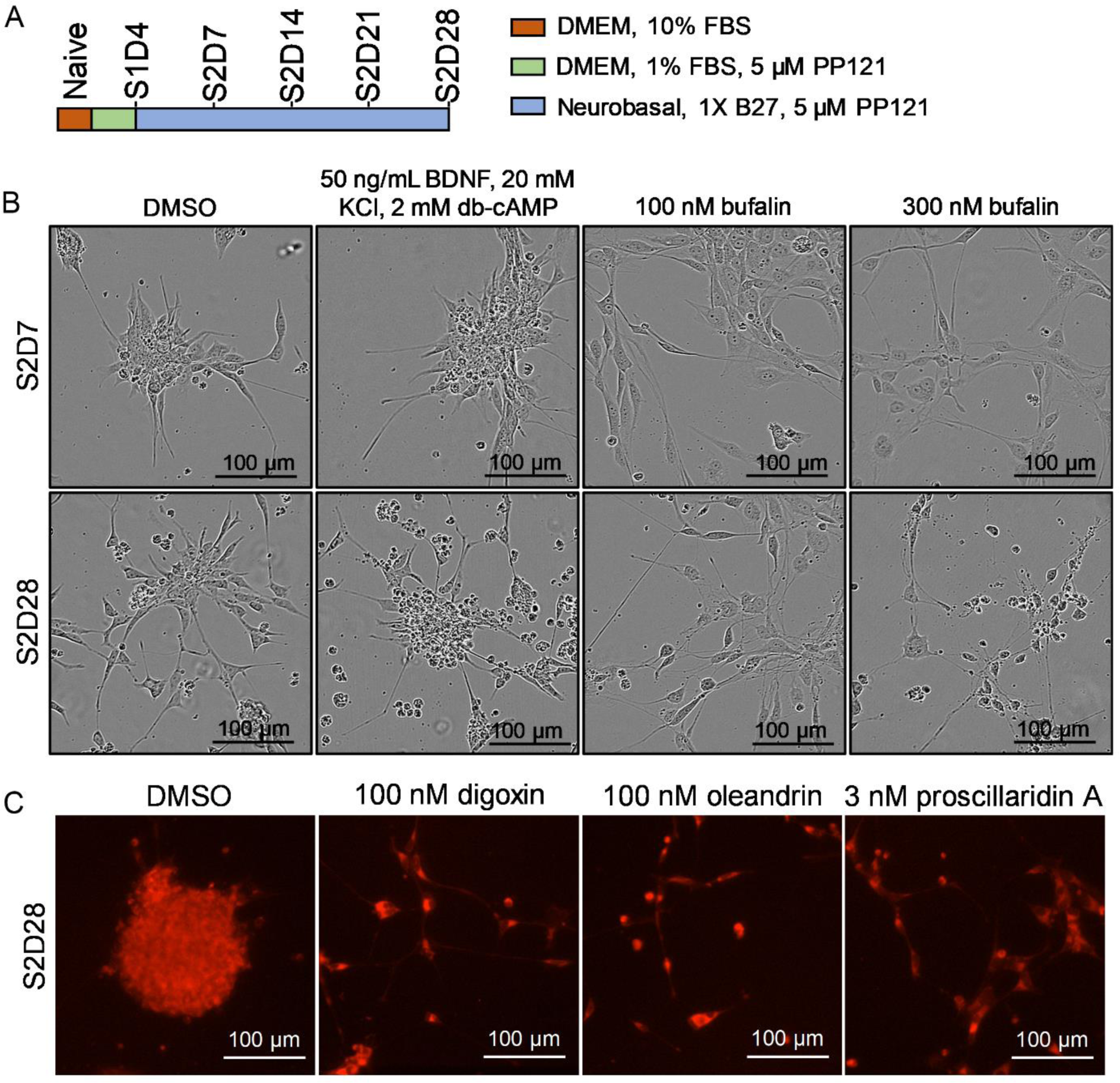
In combination with PP121, cardiac glycosides prevent clumping and promote neurite-like processes. **(A)** Differentiation protocol based on a previously established method to differentiate SH-SY-5Y cells with retinoic acid (RA). **(B)** Representative images of SK-N-AS cells at 1 week and 4 weeks in S2 medium with DMSO, BDNF/KCl/db-cAMP, and bufalin. **(C)** Representative images of SK-N-AS cells stained with a membrane TRITC dye 28 days (4 weeks) after treatment with digoxin, oleandrin, or proscillaridin A.

**Supplementary Figure 4:**
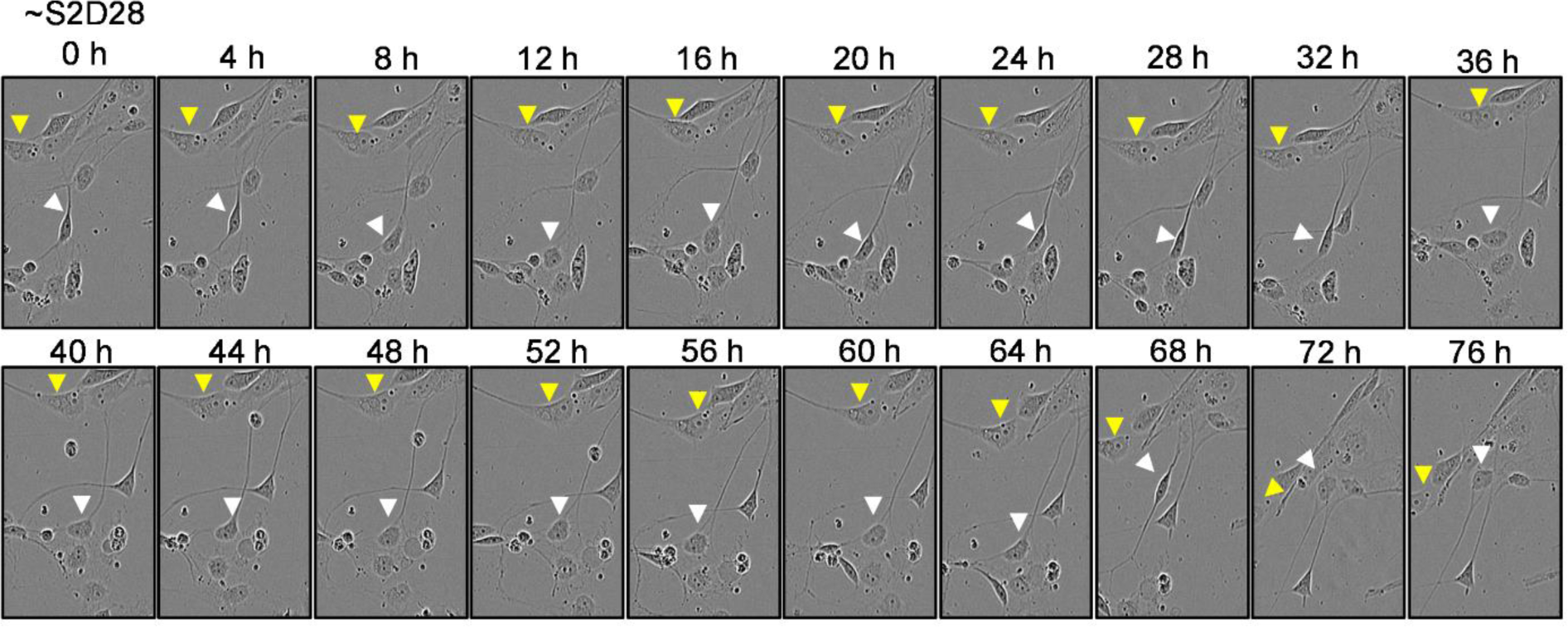
PP121/bufalin promotes long-term differentiation of SK-N-AS cells into heterogeneous cell populations. Representative images showing a heterogeneous population. The white arrow points to motile, neuron-like cells with small cell bodies and neurite-like processes; the yellow arrow points to stationary, senescent-like cells with enlarged cell bodies and no processes.

**Supplementary Figure 5:**
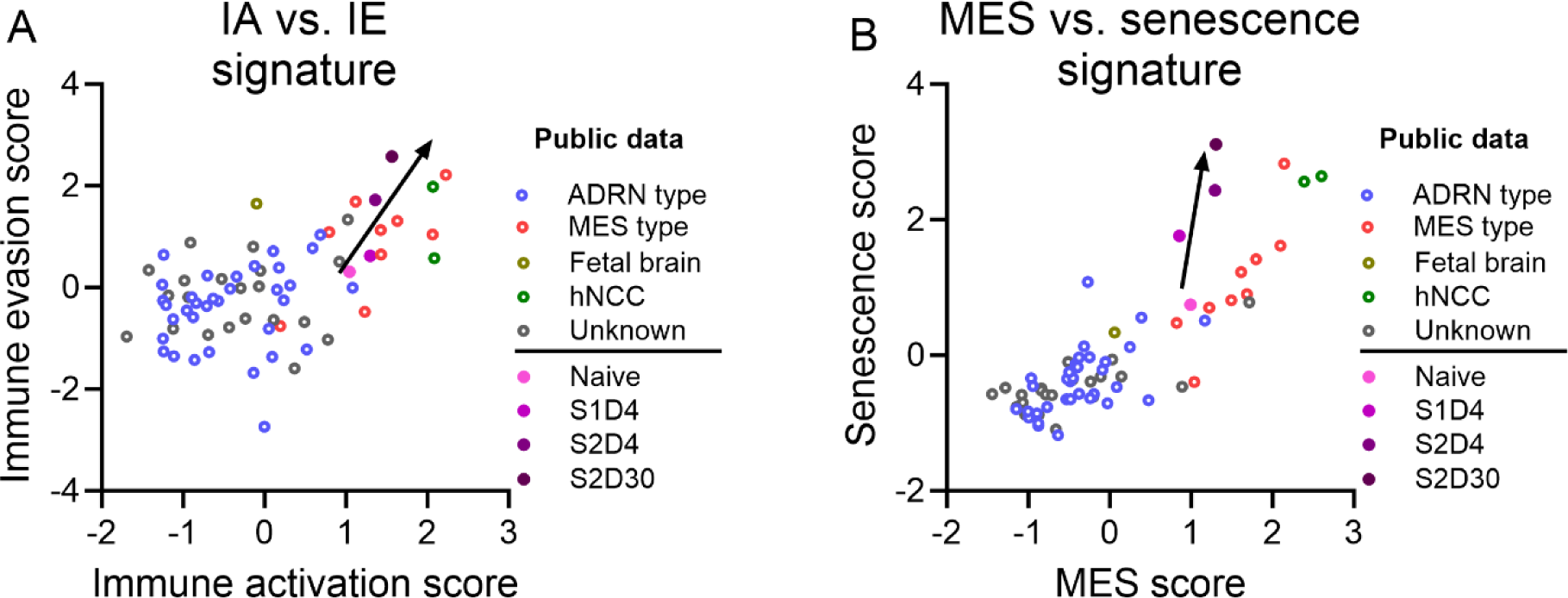
PP121/bufalin-differentiated cells became more immunogenic and senescent. **(A-B)** Differentiated SK-N-AS cells became increasingly enriched for immune activation, immune evasion, and senescence signatures (From public databases, N=67 samples; from this study, N=4 per group). Source data are provided in the source data file.

**Supplementary Figure 6:**
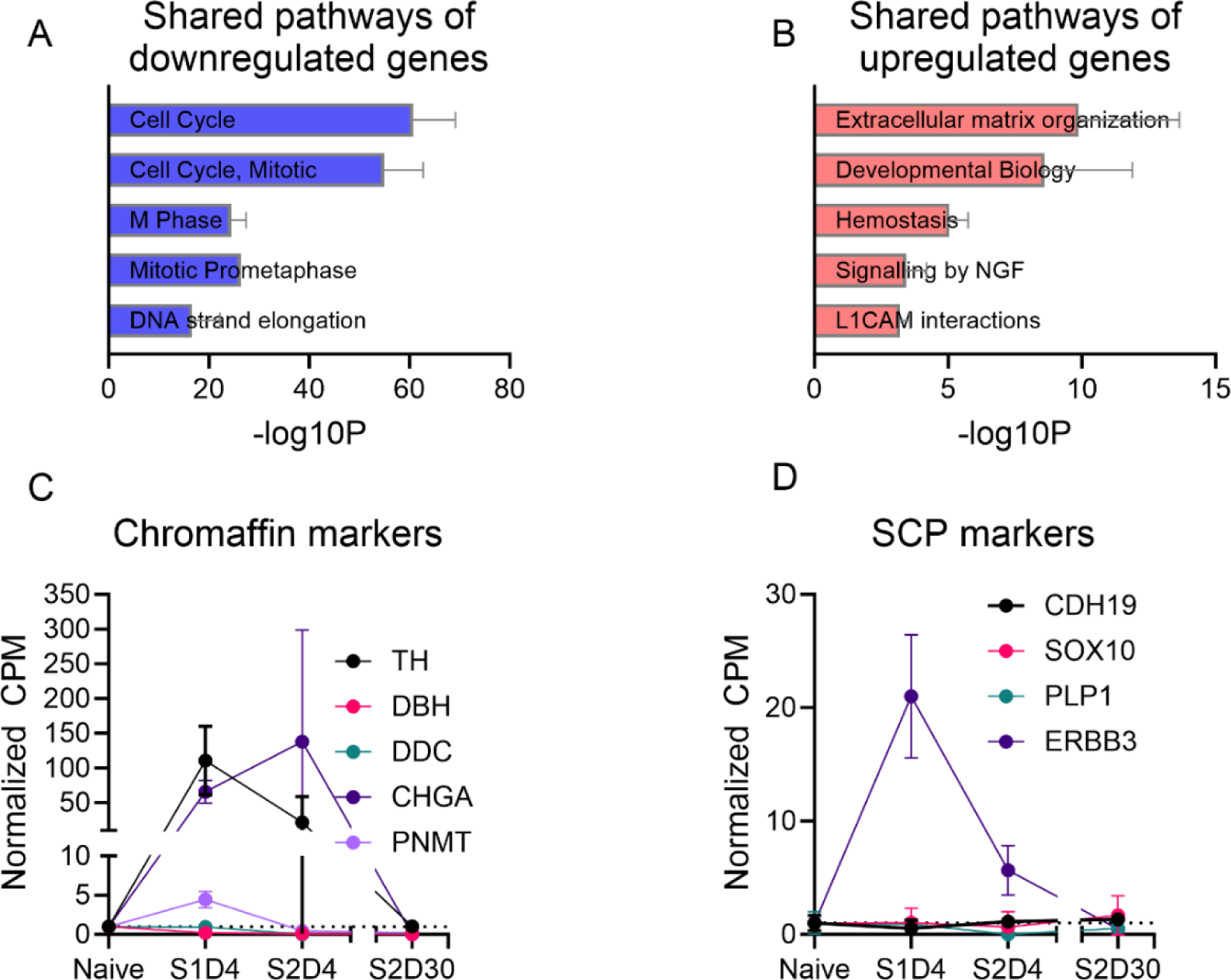
PP121/bufalin-differentiated cells are distinct from RA-differentiated cells. **(A-B)** Shared pathways of up- and downregulated genes between RA-differentiated BE2C, RA-differentiated NGP, and PP121/bufalin-differentiated SK-N-AS cells. **(C-D)** Markers associated with chromaffin cells and SCPs in naïve and differentiated SK-N-AS cells (N=4 per time point). Error bars represent S.D.. Source data are provided in the source data file.

**Supplementary Figure 7:**
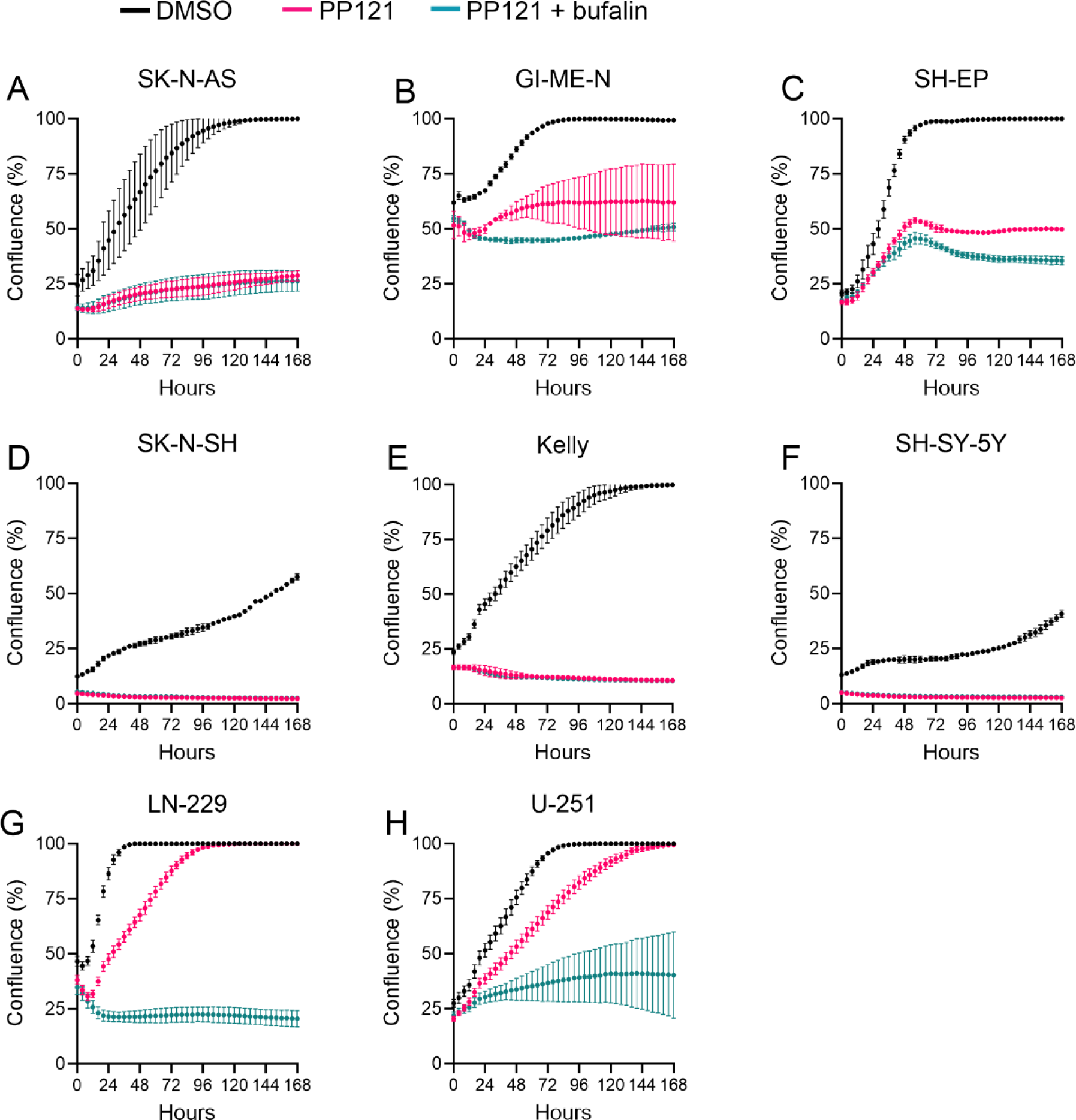
Impact of PP121/bufalin on various cell lines. **(A-C)** PP121, with or without bufalin, was sufficient to completely inhibit the proliferation of MES-type neuroblastoma cells. **(D-F)** PP121, with or without bufalin, was sufficient to completely kill ADRN-type neuroblastoma cells. **(G-H)** PP121 alone slowed down but did not completely inhibit the proliferation of glioblastoma cell lines. PP121 with bufalin completely inhibited the proliferation of glioblastoma cell lines. Error bars represent S.D.. Source data are provided in the source data files.

**Supplementary Figure 8.**
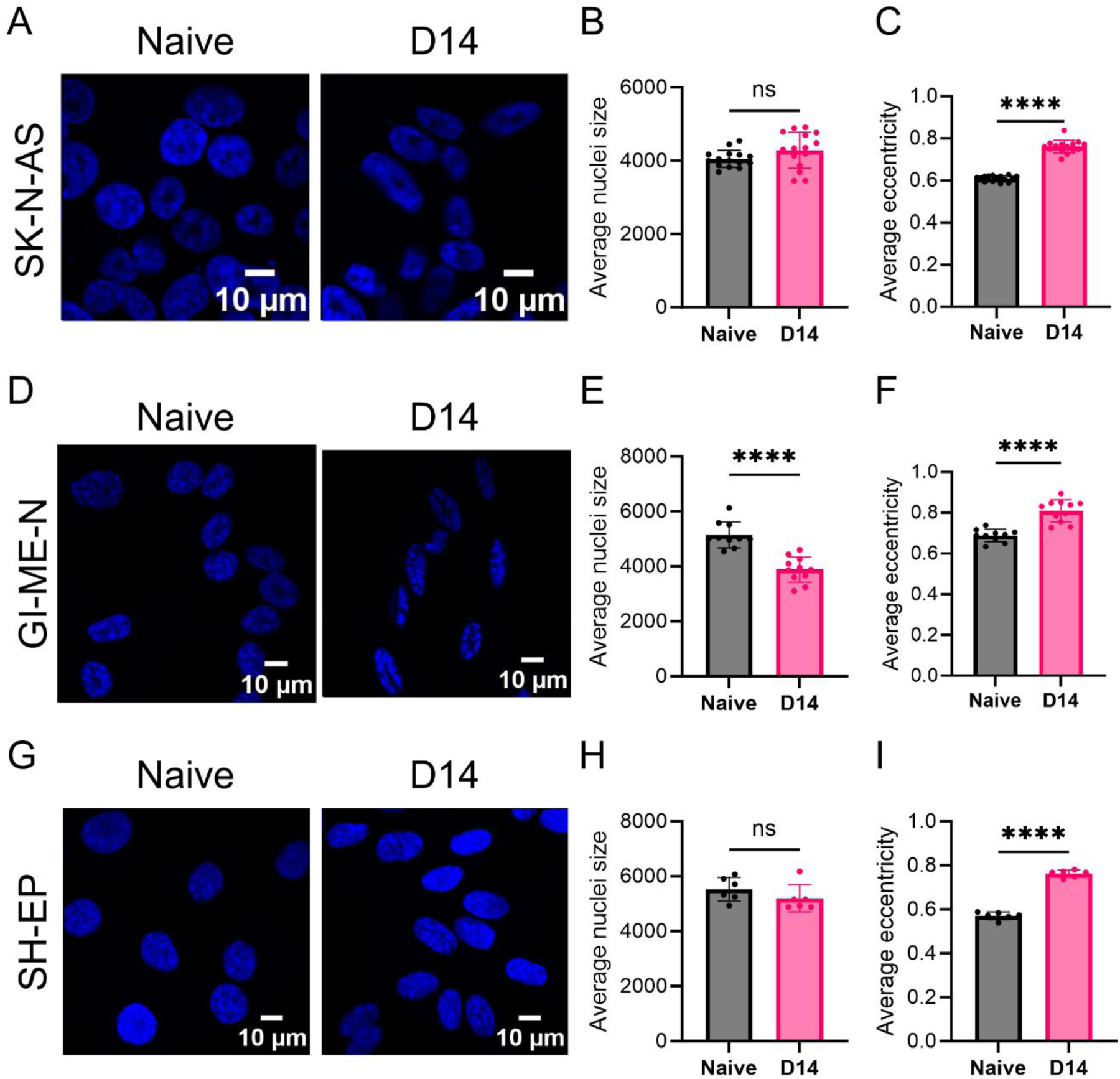
Differentiation is associated with changes in nuclear morphology. **(A-I)** In all neuroblastoma cell lines tested, differentiation was associated with an elongation of the cell nuclei as measured by the increase in nuclei eccentricity. Error bars represent S.D.. Source data are provided in the source data files.

## Supplementary Tables

**Supplementary Table 1:** List of 22 candidate compounds and their characteristics.

**Supplementary Table 2:** Confluence growth with candidate compound treatment over 4 days.

**Supplementary Table 3:** List of genes in each of the 6 clusters identified and their Z-scores.

**Supplementary Table 4:** Single-cell reference matrix for the developing adrenal medulla.

**Supplementary Table 5:** Optimized differentiation conditions for various cell lines.

**Supplementary Table 6:** Primers used in this study.

**Supplementary Table 7:** Other key resources used in this study.

**Supplementary Table 8:** CPM values of all cell lines used in this study.

## Supplementary Videos

**Supplementary Video 1:** Differentiation of SK-N-AS cells over 56 days. Cells were in Stage 1 for the first 4 days and then in Stage 2 afterward.

**Supplementary Video 2a:** SK-N-AS cells treated with vehicle (0.1% DMSO) over 7 days.

**Supplementary Video 2b:** SK-N-AS cells treated with 5 µM PP121 over 7 days.

**Supplementary Video 2c:** SK-N-AS cells treated with 5 µM PP121 + 10 nM bufalin over 7 days.

**Supplementary Video 3a:** GI-ME-N cells treated with vehicle (0.1% DMSO) over 7 days.

**Supplementary Video 3b:** GI-ME-N cells treated with 2.5 µM PP121 over 7 days.

**Supplementary Video 3c:** GI-ME-N cells treated with 2.5 µM PP121 + 10 nM bufalin over 7 days.

**Supplementary Video 4a:** SH-EP cells treated with vehicle (0.1% DMSO) over 7 days.

**Supplementary Video 4b:** SH-EP cells treated with 2.5 µM PP121 over 7 days.

**Supplementary Video 4c:** SH-EP cells treated with 2.5 µM PP121 + 10 nM bufalin over 7 days.

**Supplementary Video 5a:** LN-229 cells treated with vehicle (0.1% DMSO) over 7 days.

**Supplementary Video 5b:** LN-229 cells treated with 5 µM PP121 over 7 days.

**Supplementary Video 5c:** LN-229 cells treated with 5 µM PP121 + 30 nM bufalin over 7 days.

**Supplementary Video 6a:** U-251 cells treated with vehicle (0.1% DMSO) over 7 days.

**Supplementary Video 6b:** U-251 cells treated with 5 µM PP121 over 7 days.

**Supplementary Video 6c:** U-251 cells treated with 5 µM PP121 + 30 nM bufalin over 7 days.

**Supplementary Video 7a:** Kelly cells treated with vehicle (0.1% DMSO) over 7 days.

**Supplementary Video 7b:** Kelly cells treated with 5 µM PP121 over 7 days.

**Supplementary Video 7c:** Kelly cells treated with 5µM PP121+ 10 nM bufalin over 7 days.

**Supplementary Video 8a:** SH-SY-5Y cells treated with vehicle (0.1% DMSO) over 7 days.

**Supplementary Video 8b:** SH-SY-5Y cells treated with 5 µM PP121 over 7 days.

**Supplementary Video 8c:** SH-SY-5Y cells treated with 5µM PP121+10 nM bufalin over 7 days.

**Supplementary Video 9a:** SK-N-SH cells treated with vehicle (0.1% DMSO) over 7 days.

**Supplementary Video 9b:** SK-N-SH cells treated with 5 µM PP121 over 7 days.

**Supplementary Video 9c:** SK-N-SH cells treated with 5µM PP121+ 10 nM bufalin over 7 days.

## Notes

https://www.ncbi.nlm.nih.gov/geo/query/acc.cgi?acc=GSE270493

### Competing Interest Statement

The authors have declared no competing interest.

### Summary of Updates

(1) Expanded Figure 7. (2) Added new Supp. Fig. 1. (3) Improved discussions and fixed typos. (4) For all figures, standardized format, e.g. Arial font. (5) Added source data file and videos.

